# Deciphering the molecular landscape of human peripheral nerves: implications for diabetic peripheral neuropathy

**DOI:** 10.1101/2024.06.15.599167

**Authors:** Diana Tavares Ferreira, Breanna Q Shen, Juliet M Mwirigi, Stephanie Shiers, Ishwarya Sankaranarayanan, Miriam Kotamarti, Nikhil N Inturi, Khadijah Mazhar, Eroboghene E Ubogu, Geneva Thomas, Trapper Lalli, Dane Wukich, Theodore J Price

**Author notes:** Corresponding authors. Diana Tavares Ferreira PhD University of Texas at Dallas 800 W Campbell Rd Richardson, TX 75080 Theodore J Price PhD University of Texas at Dallas 800 W Campbell Rd Richardson, TX 75080.

## Abstract

Diabetic peripheral neuropathy (DPN) is a prevalent complication of diabetes mellitus that is caused by metabolic toxicity to peripheral axons. We aimed to gain deep mechanistic insight into the disease process using bulk and spatial RNA sequencing on tibial and sural nerves recovered from lower leg amputations in a mostly diabetic population. First, our approach comparing mixed sensory and motor tibial and purely sensory sural nerves shows key pathway differences in affected nerves, with distinct immunological features observed in sural nerves. Second, spatial transcriptomics analysis of sural nerves reveals substantial shifts in endothelial and immune cell types associated with severe axonal loss. We also find clear evidence of neuronal gene transcript changes, like *PRPH,* in nerves with axonal loss suggesting perturbed RNA transport into distal sensory axons. This motivated further investigation into neuronal mRNA localization in peripheral nerve axons generating clear evidence of robust localization of mRNAs such as *SCN9A* and *TRPV1* in human sensory axons. Our work gives new insight into the altered cellular and transcriptomic profiles in human nerves in DPN and highlights the importance of sensory axon mRNA transport as an unappreciated potential contributor to peripheral nerve degeneration.

## INTRODUCTION

Diabetes mellitus is a public health problem and diabetic peripheral neuropathy (DPN) is one of its most common and debilitating consequences, affecting an estimated 37.3 million people in the US alone (1, 2) and 463 million people worldwide (3, 4). Distal symmetric sensory polyneuropathy is the most common form of DPN, accounting for approximately 75% of DPN cases (4). The prevalence of DPN is estimated to be at least 20% in patients with type 1 diabetes (T1D) after 20 years and 50% in patients with type 2 diabetes (T2D) after 10 years (4).

DPN affects peripheral sensory nerves, and the long axons that innervate the feet are usually the first to be affected. DPN causes pain in about half of affected individuals making it the most common cause of neuropathic pain (5). Many transcriptomic, proteomic and lipidomic studies have been published using DPN rodent models to understand the underlying pathology in peripheral nerves that causes neurodegeneration and pain in diabetes (6–13). In rodent models there is clear evidence for inflammation driven by endoneurial immune cell infiltration causing increased oxidative stress and neuroimmune signaling that results in nociception and promotes axonal degeneration (7, 9, 10). Accompanying changes in the proteome (13) and lipid profile (6, 14) of peripheral nerves in rodent diabetes models are consistent with transcriptomic studies.

We are unaware of any previous studies examining transcriptomic profiles of human peripheral nerves from DPN patients, however, transcriptomic, proteomic and metabolomic studies have been conducted on human dorsal root ganglia (DRG) recovered from organ donors who died with a history of DPN pain (15, 16). These studies highlight signs of neurodegeneration (15), macrophage proliferation or infiltration (15) and altered mRNA processing (16) within the DRG associated with pain in DPN. A primary goal of our work was to use spatial and bulk RNA sequencing to understand how the peripheral tibial and sural nerves are changed in individuals with DPN. Our underlying hypothesis was that these techniques would reveal molecular insight into the extensive neurodegeneration and accompanying immune cell infiltration observed in animal studies and in previous human pathology studies. Human DRG neurons that give rise to sensory axons have a highly polarized structure with axons that extend a large distance away from the nucleus and cell body (17, 18).

Measurements from a study examining peripheral nerves in human cadavers show that the average length of the tibial nerve was 74.3 cm and that of the sural nerve was 38.4 cm (19), both of which were measured after origination from the sciatic nerve making these nerves over a meter in length in most humans. The long length of these nerves presents a challenge for maintaining the integrity of the axonal proteome because axonal transport rates from the soma do not readily account for rapid changes in protein content that can occur in axons, and these axonal transport rates are also inconsistent with maintenance of basic functions in very long axons, such as those that innervate the extremities in humans (18, 20–22). To overcome this problem peripheral axons transport mRNAs that can be translated locally to rapidly respond to external signals, and to maintain the functional integrity of axons (23). The majority of studies examining RNA transport into DRG axons have been done using rodent DRG neurons in primary cultures (24–27), although there is also strong support from *in vivo* studies for RNA transport and local translation of mRNA in DRG axons in rodents (22, 28–34). No studies have been done to assess axonal mRNAs in human peripheral axons, although we have recently demonstrated accumulation of the RNA binding protein FMRP in the central terminals of human DRG neurons (35). The secondary goal of our study was to ascertain which human peripheral nerve mRNAs originate from sensory axons using a combination of new and existing transcriptomic resources, followed by confirmation by RNA *in situ* hybridization.

In this study we used sural and tibial nerves recovered from lower leg amputation surgeries to: 1) gain insight into DPN pathogenesis and identify differences between sensory and mixed peripheral nerves and 2) begin to unravel how mRNA transport into human DRG axons may play a key role in maintaining axonal integrity in health and disease. Our findings give insight into advanced DPN pathogenesis, providing evidence for major reorganization of the Schwann cells, and fibroblasts, and alteration of endothelial and immune cell signature with disease progression. Our work also demonstrates pervasive transport of neuronal mRNAs into sensory axons with evidence that this process is guided by specific RNA binding proteins that likely bind motifs in the 3’UTRs (36–38) of these mRNAs to move them into distal axons.

## RESULTS

The study design, representative DPN sural nerve morphological photomicrographs, and the study population and sample characteristics are shown (**Figure 1**; **Suppl. File S1**). We conducted a comprehensive analysis of the tibial and sural nerves using bulk and spatial RNA-sequencing and observed that the top expressed genes were similar in these nerves (**Figure 2A, B**). Among the top expressed genes, there are ferritin light chain (*FTL*), involved in iron accumulation and apolipoprotein (*APOD*), a glia-derived apolipoprotein that has been implicated in maintaining peripheral nerve function (39). Because we had paired tibial and sural samples from the same patients (N=5), we sought to examine distinct patterns in gene expression between the sural and tibial nerves. We identified a total of 321 differential expressed genes between sural and tibial nerves (**Figure 2C**; **Suppl. File S3**). We observed that genes such as *FOSB, CXCL2, EGR1, PTGES* and *PRPH* were upregulated in sural nerves while *GABRA3, HEPACAM, CX3CR1, NEFM* and *PTGIR* were upregulated in tibial nerves (**Figure 2D**). We performed gene enrichment analysis to uncover pathways associated with these differentially expressed genes and noted that genes upregulated in sural nerve are involved primarily in non-neuronal pathways including blood vessel and vasculature development and neutrophil migration (**Figure 2E**; **Suppl. File S4)**. In contrast, genes upregulated in tibial nerve were involved in axonogenesis, axon development and glial cell migration (**Figure 2F**; **Suppl. File S5)**. This suggests that there are molecular divergences between sensory and mixed peripheral nerves in tibial and sural nerves following DPN.

**Figure 1:**
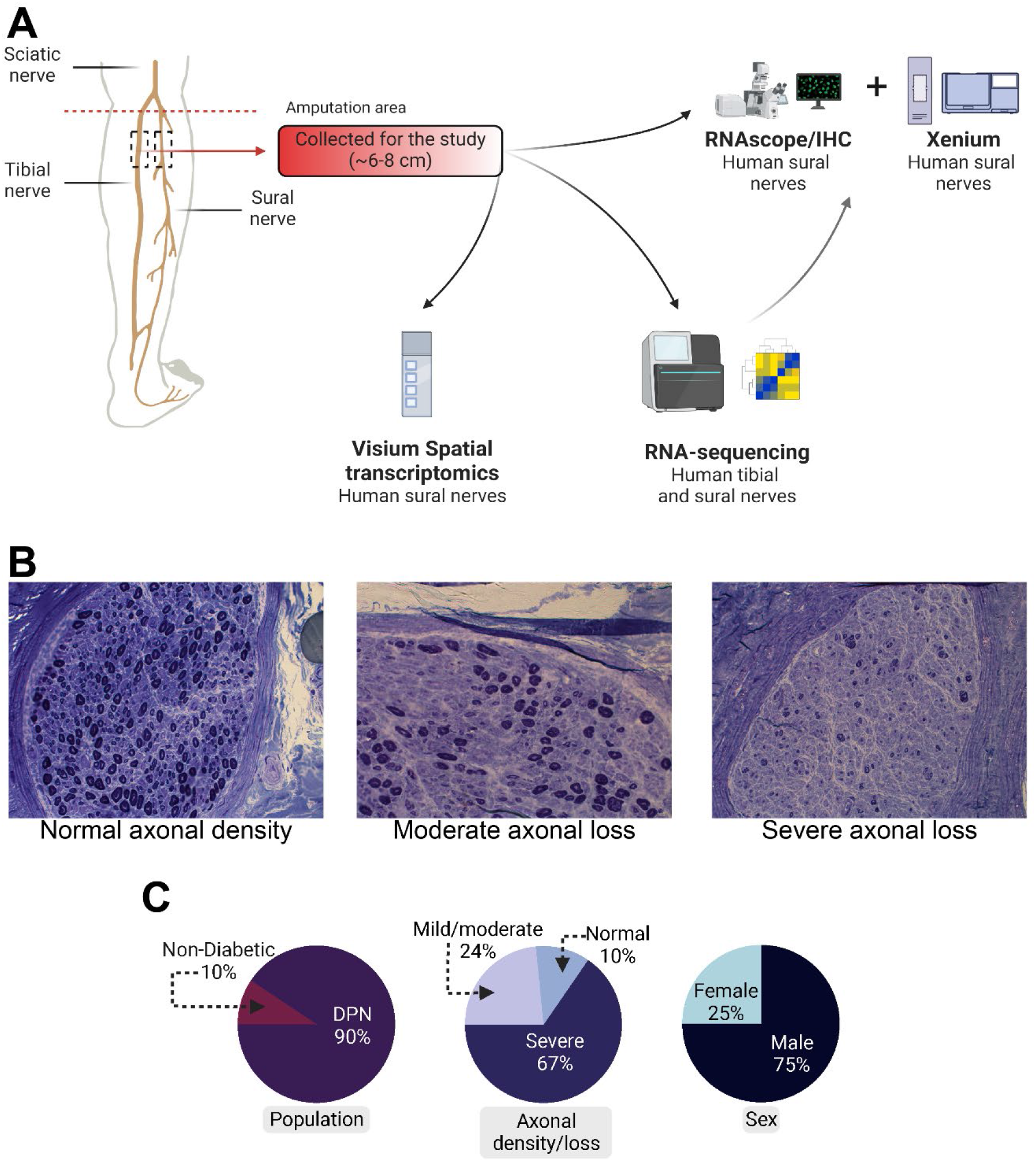
Study design. **A)** Tibial and sural nerves were recovered from amputation surgeries and processed for bulk and spatial RNA-sequencing, immunohistochemistry and in-situ hybridization (RNAscope). **B)** Representative images of axonal density in sural nerves collected from DPN patients undergoing lower extremity amputation surgeries. **C)** Population and sample description.

**Figure 2.**
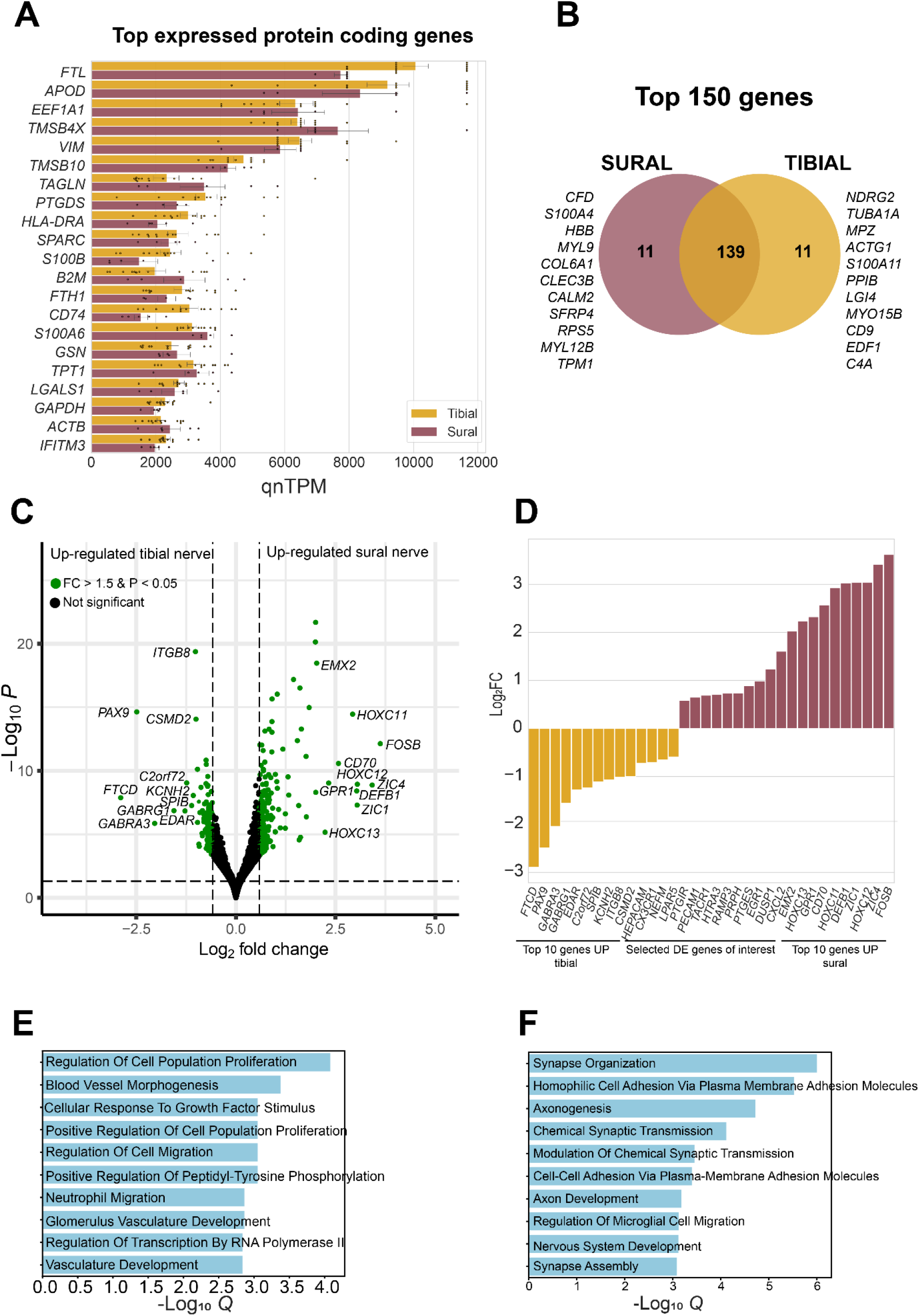
Comparisons between paired tibial and sural nerves. **A)** Top protein coding genes (excluding ribosomal protein genes). **B)** Overlap between sural and tibial top expressed genes. **C)** Differences in gene expression between sural and tibial nerve. **D)** Top differential expressed genes. Top 10 significant q-values for GO Biological Process 2023 for genes upregulated in sural **E)** and **F)** tibial nerves. Q= adjusted p-value.

Next, we leveraged visium spatial transcriptomics, which maintains spatial context, to characterize DPN sural nerves with different degrees of axonal loss (**Figure 3, Suppl. Figure 1**). Visium spatial transcriptomics uses 55 µm barcoded spots printed on specialized slides. The diameter of axons in human sural nerves ranges from 9 - 12 µm in adults (40), Schwann cells have a length ranging from 220 µm to 400 µm (with their thickness ranging between 2 and 5 µm) (41) and immune cells such as macrophages can be on average 21 µm (42) and T cells vary between 8–10 µm in diameter (43). Therefore, we first wanted to characterize the enriched cell types in a given barcoded spot. We used a sample with normal axonal density as a reference to identify the major cell types present in human sural nerves. We identified 5 major cell types in human sural nerve (**Figure 3A-C**): fibroblasts (*FBLN1, TNXB*), endothelial cells (*AQP1*), Schwann cells (*SOX10, MPZ*), immune cells (*PLCG2, CD74*) and adipocytes (*PLIN1*). One of the identified enriched cell types expressed SRY-Box Transcription Factor 10 (*SOX10*). We hypothesized that SOX10 may be a marker for a subtype of Schwann cells. Using RNAscope and IHC (two different antibodies), we found that *SOX10* mRNA transcripts were highly colocalized with SOX10 protein (**Suppl. Figure 2).** Additionally, *SOX10* mRNA puncta were particularly localized in Schwann cells surrounding nerve fibers that colocalized with DAPI, which stains cell nuclei. Schwann cells, the glial cells in peripheral nerves, are key to maintaining axonal homeostasis, encasing them in myelin, supporting them through release of neurotrophins such as Nerve Growth Factor (NGF) (44), providing rationale to further evaluate how this cell type changed in nerves from DPN patients with moderate and severe axonal loss.

**Figure 3.**
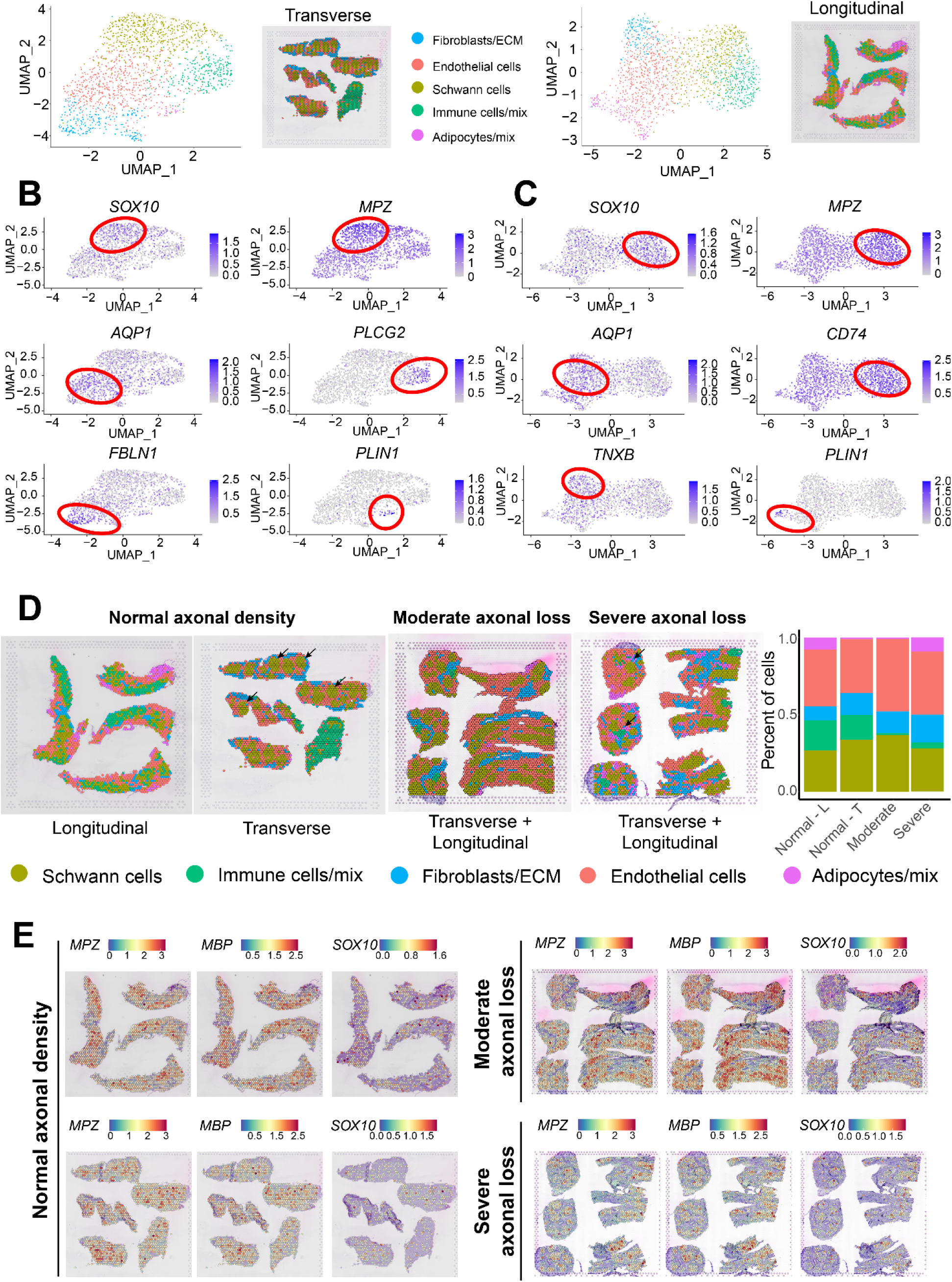
Spatial analysis of human sural nerves. **A)** Using the normal axonal density sample as a reference, we identified 5 major cell types enriched in sural nerves. Markers used to identify cell types in **(B)** transverse and **(C)** longitudinal sural nerve sections. **D)** We identify a spatial shift in different cell types across different levels of axonal density/loss. **E)** We observe a decrease in Schwann cell and myelin genes in sural nerves with severe axonal loss. L-longitudinal; T -transverse.

We applied the same approach to samples with moderate and severe axonal loss and observed changes in cell types associated with axonal loss severity (**Figure 3D**). We observed a reduction in the relative proportion of Schwann cells within sural nerves with a relative increase in endothelial cells, fibroblasts and immune cells in the sural nerve with severe axonal loss (**Figure 3D)**. At the gene expression level, we observed that Schwann cells and myelin-associated genes were markedly decreased, particularly in the sample with severe axonal loss (**Figure 3E**). We then conducted cell-cell interaction analysis using the Cellchat package (45) with focus on sural nerves with moderate and severe axonal loss and observed multiple cell-to-cell interactions occurring in advanced DPN (**Figure 4**). Our analysis showed that Schwann cells, fibroblasts and immune cells had the highest number of interactions in advanced DPN (**Figure 4-A**,**C**). With severe axonal loss, we saw an increase in interactions with adipocytes, suggesting that these cells are associated with DPN progression. Collagen was the top signaling pathway for both moderate and severe axonal loss samples (**Figure 4-E**,**F**). Collagens are important components of the extracellular matrix and of the connective layers in the peripheral nerve and our results suggest that changes occurring in these pathways may be involved in the nerve degeneration following DPN. The ligand-receptor pairs involved in these pathways are listed in **suppl. Figure 3.** We also analyzed possible interactions between ligands expressed by these enriched cell groups and receptors that are present in human DRG neurons **(Suppl. Figure 4)**. We identified several interactions that are driven by ligands such as Amyloid Beta Precursor Protein (*APP*) which is involved in fiber organization and neuron remodeling. This suggests that the ligands expressed in sural nerves are likely involved in axonal degeneration and regeneration.

**Figure 4.**
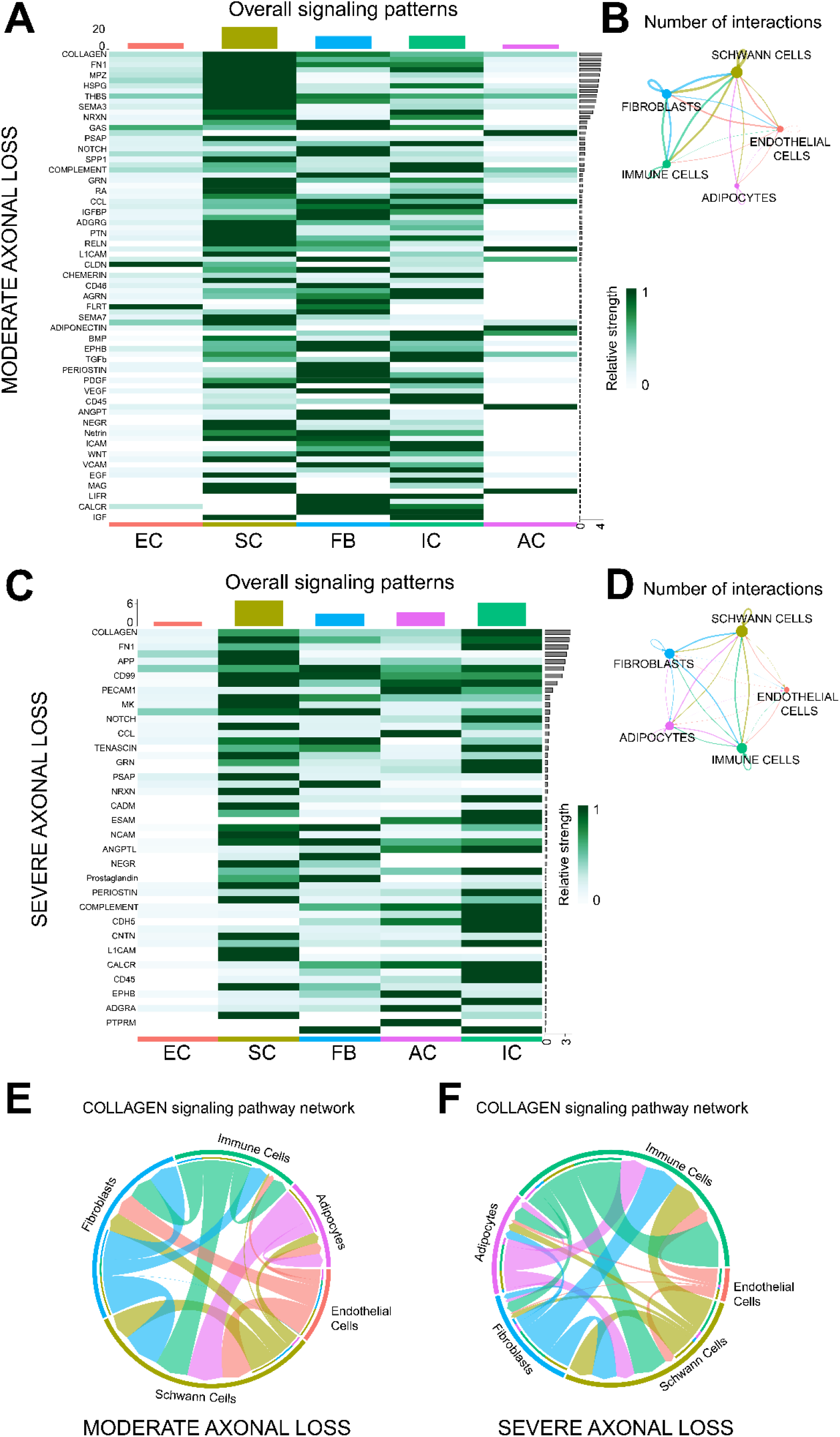
Overview of cell-cell interaction analysis in sural nerves with moderate and severe axonal loss. **A, C)** Heatmap showing the overall contribution of signaling pathways in the enriched cell types in a sample with moderate axonal loss **(A)** and in a sample with severe axonal loss **(C).** Values are scaled per row. **B, D)** Circle plot displaying all the interactions in each cell group in moderate **(B)** and severe **(D)** axonal loss. The thickness of each line illustrates the quantity of interactions between two distinct cell types, whereas the size of each dot signifies the overall quantity of interactions that involve that particular cell type. **E, F)** Chord diagram displaying the interactions within the collagen signaling pathway network in moderate (**E)** and severe **(F)** axonal loss nerves. Analysis and plots generated using Cellchat package (45). EC: Endothelial cells, SC: Schwann cells; FB: Fibroblasts; IC: Immune cells; AC: Adipocytes.

Next, we sought to confirm our findings using an independent approach choosing cell type deconvolution to examine cell proportions within barcodes. We utilized a reference-free cell type deconvolution method, STdeconvolve (46), and characterized moderate and severe axonal loss DPN sural nerves (the visium frame for these samples contained both transverse and longitudinal sections). We observed similar results to our enriched cell type analysis and demonstrated that cell types drastically change with axonal loss severity (**Suppl. Figure 5**).

Using cell type deconvolution, we identified gene expression markers for axons, myelinating Schwann cells, smooth muscle cells, perineurium and other connective layers in both nerves with moderate and severe axonal loss. Non-myelinating Schwann cells were present only with moderate axonal loss. Additionally, mitochondrial gene expression was observed in DPN with several axonal loss, implying activation of oxidative stress pathways with advanced disease.

Guided by considerable changes in peripheral cell types associated with changes in axonal density, we compared the transcriptome of DPN sural nerves with moderate and severe axonal loss using bulk RNA-sequencing data (**Figure 5**; **Suppl. File S6)**. Nerves with moderate axonal loss were significantly enriched in genes such as Baculoviral IAP Repeat Containing 7 (*BIRC7*) which is a family member of inhibitors of apoptosis (47), ALK receptor tyrosine kinase (*ALK*) and myelin associated glycoprotein (*MAG*). Nerves with severe axonal loss showed an increase in genes such as peripherin (*PRPH*), calveolin-1 (*CAV1*) and Collagen Type XXV Alpha 1 Chain (*COL25A1)* (**Figure 5A,B**). Using imaging-based spatial transcriptomics (Xenium by 10x Genomics) and their pre-designed brain panel, we verified that *ALK* is expressed mostly in immune cells, *MAG* is expressed in Schwann cells. *COL25A1* and *CAV1* are expressed in fibroblasts; *CAV1* is also expressed in endothelial cells (**Suppl. Figures 6)**. These data suggest that multiple peripheral cell types (immune cells, fibroblasts, Schwann cells and endothelial cells) are altered and involved in the pathogenesis of advanced DPN with moderate and several axonal loss. Our gene enrichment analysis showed that differentially expressed genes increased in severe axonal loss were involved in metabolic process, wound healing and signaling pathways such as signal transducers and activators of transcription (STAT) and transforming growth factor beta (TGFB) receptor pathways (**Figure 5C**; **Suppl. File S7**).

**Figure 5.**
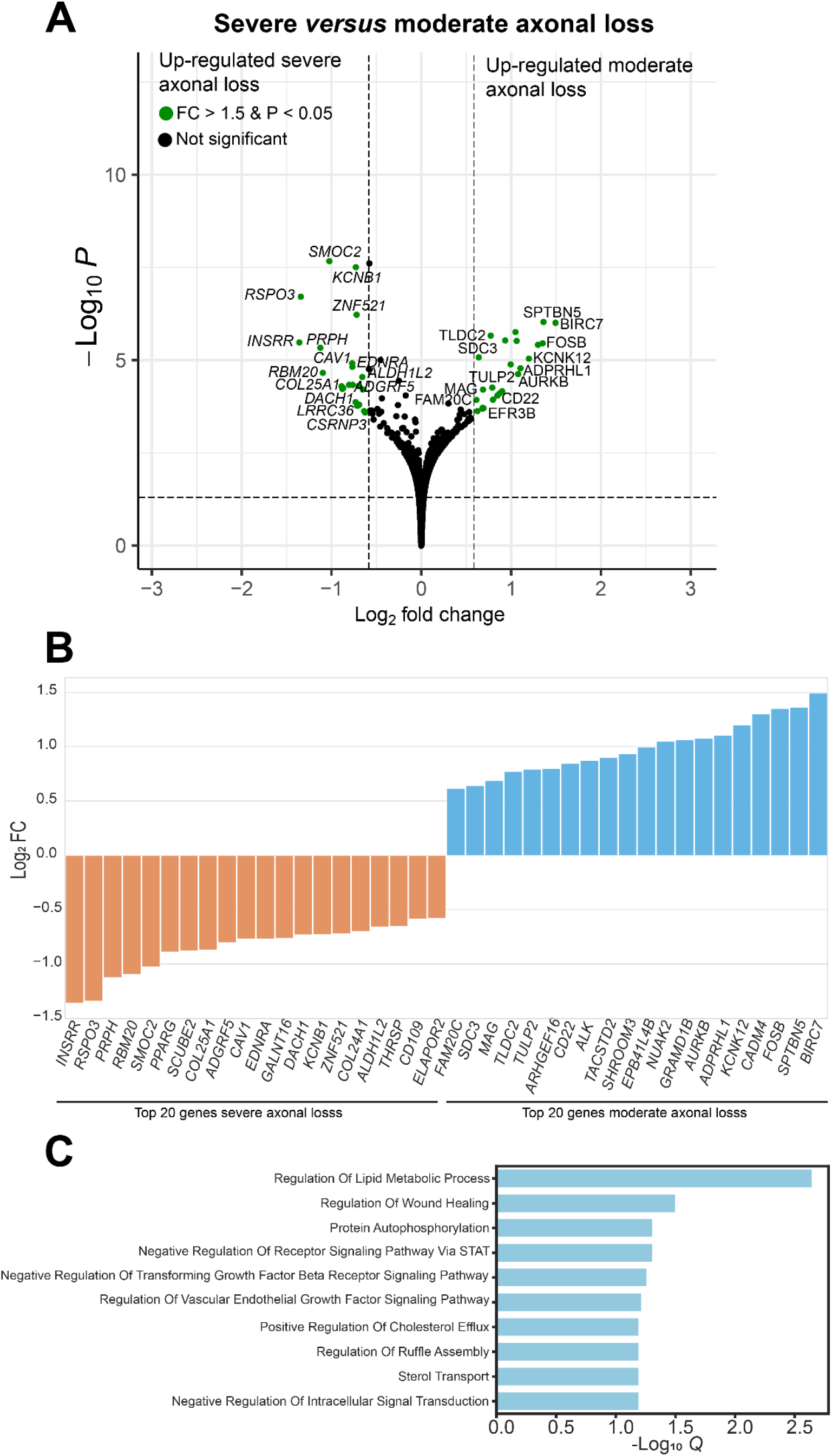
Comparison between sural nerves with severe and moderate axonal loss. **A)** Differential gene expression between sural nerves with severe versus moderate axonal loss. **B)** Top genes differential expressed. **C)** Gene enrichment analysis shows the top pathways affected by the differential expressed genes. Q= adjusted p-value.

Peripherin is an intermediate filament important for neuronal function and the differences we identified at the mRNA level between sural nerves with moderate and severe axonal loss led us to investigate axonal mRNA localization. This is an unstudied area of human sensory neuron research; however, previous rodent studies have shown that local translation is crucial for the maintenance of axonal homeostasis and regeneration (26, 48, 49). As axonal degeneration is commonly observed in peripheral neuropathies, including DPN, we examined the expression of neuronal markers in human peripheral nerves. Using bulk RNA-sequencing experiments, we detected neuronal markers such as transient receptor potential vanilloid 1 (*TRPV1)* and sodium voltage-gated channel alpha subunit 9 (*SCN9A)* in human tibial and sural nerves (**Figure 6A**). We also conducted a comprehensive meta-analysis of previously published studies (**Suppl. Figure 7A**) and identified a distinct set of genes that appear to be exclusively in sensory axons, representing putative axonal mRNAs (**Suppl. Figure 7B**). To strengthen our hypothesis that these genes are axonal and originate from the soma, we show their expression in human DRG neurons (**Suppl. Figure 7C)**. Next, we performed RNAscope *in situ* hybridization on human sural and sciatic nerves for the following neuronal markers: *SCN9A*, *TRPV1*, *PRPH*, neurotrophic receptor tyrosine kinase 1 (*NTRK1*) and sodium voltage-gated channel alpha subunit 10 (*SCN10A*) followed by immunohistochemistry (IHC) to label cell nuclei (DAPI) and axons (peripherin) to provide direct evidence for RNA localization for these genes in human peripheral nerve axons.

**Figure 6.**
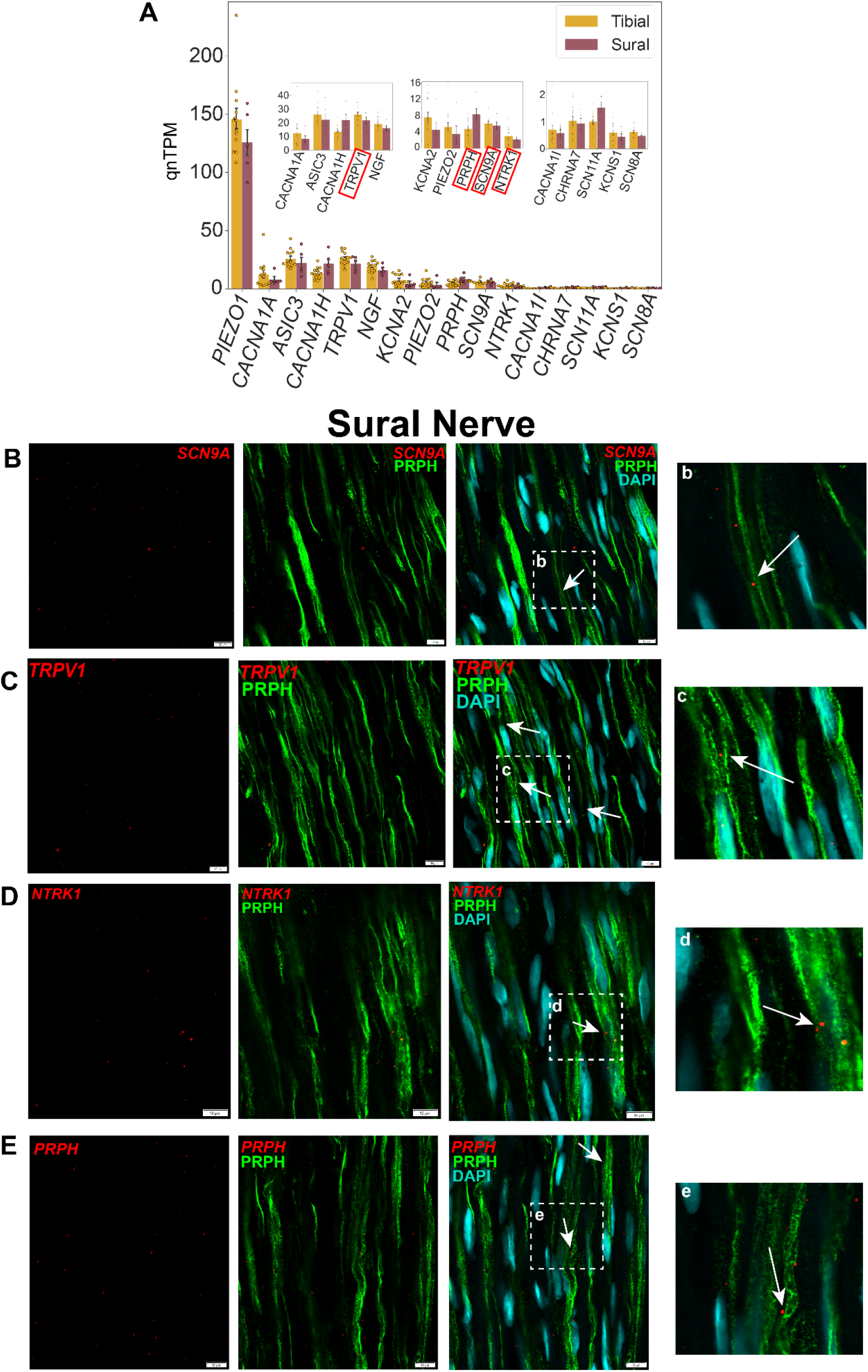
Gene expression in human peripheral nerves. **A)** We performed bulk RNA-seq of human tibial and sural nerves and detected the presence of several mRNAs involved in sensory processing such as *TRPV1* and *SCN9A*. mRNA puncta of *SCN9A* **(B)**, *TRPV1* **(C)**, *NTRK1* **(D)**, *PRPH* **(E)** in red are colocalized with peripherin (PRPH, green), which labels nerve fibers. Arrows point to areas where mRNA puncta do not overlap with DAPI (cyan), suggesting that it is axonal specific staining. Insets show zoomed-in images. Scale bars=10 µm.

*SCN9A*, which encodes Nav1.7, colocalized with peripherin in sural and sciatic nerves (**Figure 6B** and **Suppl. Figure 8A**, respectively). Overlap of *SCN9A* puncta with peripherin was present even in areas where there was no DAPI staining, suggesting that these mRNAs exist in both the nuclei of axon-supporting cells, as well as in the nerve axons. While we did not quantify RNA expression levels, the number of *SCN9A* puncta was comparable between sural and sciatic nerves. Colocalization with peripherin, in areas without DAPI staining, was also seen in sural and sciatic nerve for mRNA puncta of *TRPV1* (**Figure 6C** and **Suppl. Figure 8B**, respectively), which encodes the transient receptor potential vanilloid 1 nonselective cation channel receptor, *NTRK1* (**Figure 6D** and **Suppl. Figure 8**C, respectively), which encodes the TrkA receptor tyrosine kinase, and *PRPH* (**Figure 6E** and **Suppl. Figure 8D**, respectively).

While *SCN10A* (which encodes Nav1.8) was not detected with tibial and sural nerve bulk-RNA sequencing, Nav1.8 has an important role in nociception, so we decided to further investigate *SCN10A* axonal expression with a more sensitive technique, guided by previous reports of this mRNA localizing to rat sensory axons originating from DRG (28, 50). We observed only a few mRNA puncta colocalized with peripherin; however, this observation suggests transport for this mRNA in human peripheral nerve axons as well (**Suppl. Figure 9**). The difference in sensitivity between bulk-RNA sequencing and RNAscope may contribute to the detection of a few *SCN10A* transcripts with the latter approach. In line with our bulk RNA-sequencing data, our qualitative assessment of mRNA puncta suggests that *SCN9A* is more highly expressed in sural and sciatic nerves than the other mRNAs. Additionally, using high-resolution imaging-based Xenium approach, we were able to detect neuronal markers such as parvalbumin (*PVALB*), tachykinin precursor 1 (*TAC1*), and transient receptor potential cation channel subfamily C member 5 (*TRPC5*) in peripherin-labelled human peripheral axons in longitudinal and transverse sections (**Suppl. Figure 10**). We also examined markers of neuronal subtypes previously identified in the human DRG (51) and we identified mRNA expression for 96 out of 126 unique gene markers (76.19%) in the human tibial and sural nerve bulk RNA-seq data (**Suppl. Figure 11)**. Identification of specific neuronal subpopulation markers can provide important information for the identification of specific subtypes potentially affected in DPN.

Along with cis-acting elements (or motifs) usually found in the 3’ untranslated region of mRNAs (52, 53), RNA-binding proteins (RBPs) are required for RNA transport (54) and this process can be species-specific (55–57). After demonstrating that neuronal genes *SCN9A, SCN10A, TRPV1, PRPH,* and *NTRK1* are present in peripheral nerve axons, we sought to identify which RBPs were present in human peripheral nerves. We performed the Somascan proteomic assay and detected 1890 RBPs, including 40 RBPs with a role in RNA transport (based on RBPs database RBP2GO (58)), in paired L4 DRG and peripheral nerves from the same organ donors (N=6, **Figure 7, Suppl. File S8**). RBPs such as Fragile X mental retardation protein (FMRP), which is encoded by the *FMR1* gene, is known to play an important role in RNA transport (59, 60). We found that FMRP was robustly detected in the DRG and sciatic nerve samples from organ donors, including those with a pain history (**Figure 7B)**. Using IHC, we validated that FMRP protein was present within human sural nerves showing that the protein localizes to distal sensory axons (**Figure 7C**).

**Figure 7.**
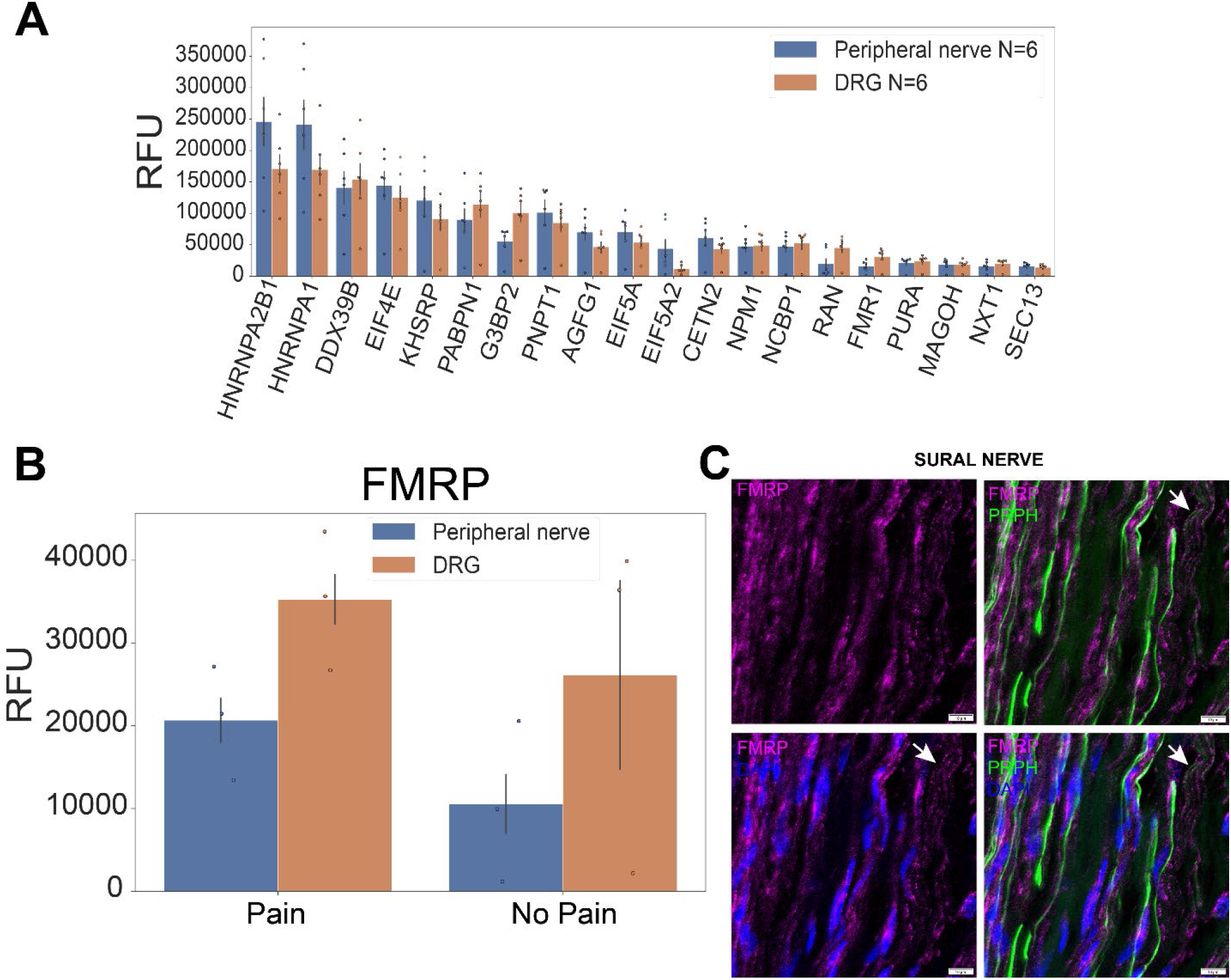
RNA-binding proteins (RBPs) in the human peripheral nervous system. **A)** Top RBPs associated with RNA transport in human DRG and sciatic nerve. **B)** Changes in FMRP in peripheral nerve and DRG from donors with and without pain history. **C)** We detect the presence of FMRP in human sural axons using immunohistochemistry. Arrows point to area where FMRP is not co-localized with DAPI staining. Scale bar= 10 µm. Suppl. Figures and Tables are available in a separate pdf document. Suppl. Files are provided in separate Excel files.

## DISCUSSION

Peripheral nerves are responsible for signal transduction to and from the central nervous system (CNS). In humans, sensory axons extend for a large distance and maintaining axonal integrity is crucial for normal function. Axonal damage such as that observed in DPN, typically affects sensory axons in a length-dependent manner, highlighting the importance of uncovering the pathogenic cellular and molecular alterations, including changes in axonal mRNA transport. Previous human and rodent studies implicated that inflammatory mediators released by different cell types play an important role in the development and progression of DPN (61). Additionally, studies on sural nerve biopsies found alterations in immune response, calcium signaling, and axon guidance (62–64). In this study, we characterized human tibial and sural nerves using a multi-omics approach. We found distinct pathways enriched in sural (sensory) and tibial (mixed sensory and motor) nerves following DPN. In sural nerves, we observed an enrichment in non-neuronal pathways including vasculature development and neutrophil migration. These findings support the notion that inflammatory processes and vascular alterations are pathologically important in DPN, affecting nerve function indirectly through microenvironment changes (65). In tibial nerves, the enriched pathways consisted of axonal biology-related terms. This suggests that the mechanisms of injury and repair may vary significantly between different peripheral nerves in DPN. Patients with DPN often describe sensory symptoms such as pain, tingling or numbness but motor symptoms like weakness and loss of coordination are also reported, particularly in patients with distal symmetric sensorimotor polyneuropathy (66). Additionally, nerve conduction studies have demonstrated that alterations in the amplitude of motor nerve fibers generally occur after those observed in sensory nerve fibers (67). Unlike the sural nerve, which is a purely sensory nerve, the tibial nerve contains a mix of motor and sensory axons, so it is feasible that this difference contributes to transcript differences seen in our paired samples.

An important component of neuronal homeostasis on neuroimmune and neuro-glial interactions (68, 69) and inflammatory mediators released by different cell types play an important role in the development and progression of DPN as previously reported (61). Schwann cells are essential for the structure and function of peripheral nerves and contribute to DPN pathogenesis via oxidative stress, endoplasmic reticulum stress and inflammation (70–72). Additionally, their impaired ability to produce neurotrophic factors crucial for nerve health, coupled with dyslipidemia associated with diabetes that alter the composition of myelin sheaths, leads to axonal degeneration and aberrant signal transmission (70). Using our spatial RNA-sequencing data, we observed changes in Schwann cells, immune cells, fibroblasts, and endothelial cells associated with axonal loss severity in DPN. First, using our cell enrichment analysis approach we observed a decrease in Schwann cells and their gene markers in DPN with severe axonal loss. Second, using our cell type deconvolution approach we found that non-myelinating Schwann cells were the most affected, possibly as consequence of unmyelinated axon loss. Both myelinating and non-myelinating Schwann cells are affected by DPN. Myelinating Schwann cells are primarily responsible for forming the myelin sheath around axons, but in DPN, they often exhibit metabolic dysregulation due to the effects of hyperglycemia and disrupted insulin signaling, leading to myelin sheath abnormalities and nerve function impairment (73). Non-myelinating Schwann cells, which enwrap multiple unmyelinated axons, are also crucial for the maintenance and integrity of peripheral nerves (74); however, in DPN, these cells are more susceptible to the toxic effects of hyperglycemia, impairing their ability to support unmyelinated nerve axons that are responsible for nociception (75). Previous research has shown that non-myelinating Schwann cells create wide signaling networks with immature Schwann cells and macrophages in an attempt to protect nerve function (76).

Accordingly, our interactome analysis shows that Schwann cells establish interactions with immune cells. In DPN with severe axonal loss, we observed a substantial loss in Schwann cells and particularly non-myelinating Schwann cells that is also reflected in the reduced number of interactions. Instead, there is a relative increase in interactions involving immune cells and adipocytes. Our interactome analysis also revealed that cell-cell signaling pathways occur between the enriched cell groups, particularly related to collagen and extracellular matrix. The extracellular matrix and connective tissue layers of the peripheral nerves (epineurium, perineurium, endoneurium) play an important role in preserving nerve integrity and in the regenerative response to injury. A previous study showed that collagen V and VI were increased in DPN patient sural nerve biopsies and may contribute to the progression and limited regenerative response in DPN (77). This suggests that targeting these pathways early could offer an opportunity to revert to a pro-regenerative phenotype. We also observed that the top ligands expressed in the nerves can interact with receptors in DRG neurons. Interestingly, ligands such as *APP* and *C3* are involved in axonal reorganization and remodeling, which suggests that peripheral nerve cells can release mediators that directly affect axons. Complement C3 has been associated with increased risk of neuropathy (78), suggesting that it may have a pro-inflammatory role that limits the nerve regeneration capacity. Similarly, APP has been associated with neuronal death in the CNS (79, 80) and may contribute to axonal degeneration in DPN.

Due to shifts in peripheral nerve cell types identified between DPN sural nerves with moderate and severe axonal loss using our spatial approach, we set to investigate the molecular changes using bulk RNA-sequencing. Peripherin is an intermediate neurofilament and a structural component of axons. It has been identified as a marker of axonal damage (81) and functions in neurite stability and axonal transport (82). Peripherin antibodies were present in patients with type I diabetes, and a reduction in peripherin expression was observed to accompany hyperalgesia in a rat streptozotocin-induced type I diabetes model (82, 83). In our study, we observed an increase of peripherin mRNA in DPN sural nerves with severe axonal loss, suggesting that *PRPH* can also be a marker of axonal loss in human sural nerves. A previous study found that diabetic mice lacking neurofilaments experienced significant conduction velocity slowing and decreased nerve action potential amplitude, unlike those with normal neurofilaments who showed only mild neuropathy, irrespective of hyperglycemia levels (84). This indicates that neurofilaments aid axons resist diabetic damage. We also observed that caveolin-1 (*Cav1*) is upregulated in DPN sural nerves with severe axonal loss. Previous studies in mice had shown that absence of Cav1 correlated with increased DPN severity, including motor and sensory nerve conduction velocities, and mechanical or thermal sensitivity (85) and that low levels of Cav1 contribute to demyelination (86). Cav1 has also been associated with pain development (87) as well as anti-inflammation and neuroprotection (88). These observations suggest that there are regenerative pathways present in peripheral nerves that can be targeted to facilitate recovery in DPN.

In this study, we also characterized the expression of neuronal mRNAs *SCN9A, SCN10A, TRPV1, NTRK1,* and *PRPH* with RNAscope *in situ* hybridization in human sural and sciatic nerves. We demonstrated axonal localization of these mRNAs, consistent with our RNA sequencing data. We qualitatively observed similar levels of each mRNA in both sciatic nerves isolated from organ donors and sural nerves from surgeries. Axonal mRNA transport is important for development, regeneration and response to injury or endogenous molecule signaling (89). During pathfinding in normal development, axons respond to neurotropic cues through local protein synthesis (90). After tissue damage, translation can be initiated in DRG neurons and their axons, resulting in sensory neuron sensitization (91). Local mRNA translation can be advantageous by allowing for faster responses to stimuli than axoplasmic transport of proteins, which can take hours or days to traffic proteins to peripheral axon segments (92).

Following axonal translation, locally synthesized mRNAs can also be retrogradely transported to neuronal soma and alter nuclear transcription, raising another mechanism through which peripheral protein synthesis can mediate nociceptive response (30, 60, 93). While rodent sensory axon transport and local protein synthesis had been shown for some mRNAs relevant to nociception, including Nav1.8 (34, 91), there is still much work to be done characterizing which mRNAs are translated downstream of nociceptive input, particularly in human peripheral nerves. In this study, we have identified for the first time mRNAs that are localized to the human sensory axons. Revealing which RNAs are transported into axons, and therefore can potentially be translated locally, could result in novel therapeutic targets for pain and axonal regeneration in DPN and other peripheral neuropathies.

Directing mRNAs to specific subcellular sites requires three major components: 1) cis-acting elements within the mRNA, most frequently found in the 3′ UnTranslated Region (UTR); 2) RNA-binding proteins (RBPs) that can recognize and bind to the cis-acting elements in a sequence-specific manner; and 3) the resulting ribonucleoprotein (RNP) complex that can, then, be linked directly to motor proteins or hitchhike to vesicles such as lysosomes and mitochondria for transport to a specific subcellular region (94). Our analysis revealed that several RBPs are present in peripheral nerves, including fragile X mental retardation protein (FMRP). FMRP is a regulatory RBP that controls mRNA transport and local translation with known roles in neurodevelopment and synapse function (59, 60). Previous studies have shown that disruption of RBPs involved in mRNA axonal transport can consequently lead to the loss of axonal integrity (95). This can result in neuronal degeneration and cause neurological diseases such as fragile X syndrome (FXS) (96), amyotrophic lateral sclerosis (ALS) (97), spinal muscular atrophy (SMA) (98) and peripheral neuropathy (99). Additionally, peripheral neuropathy is a common and significant feature of Fragile X-associated Tremor/Ataxia Syndrome (FXTAS), involving damage to the peripheral nerve (100–102). FXTAS is a neurodegenerative disorder in individuals with the Fragile X Mental Retardation 1 *(FMR1)* premutation that leads to cognitive impairment, tremors and neuropathy. Males carrying the *FMR1* premutation showed a loss of distal reflexes and a reduction in vibratory perception, and a strong correlation was identified between CGG repeat length and total neuropathy score in both males and females (102). The X-inactivation ratio, which determines the relative expression of normal versus premutation alleles, is thought to influence the prevalence and severity of symptoms in females (103). In line with these studies, our proteomics data show for the first time the potential involvement of FMRP in DPN, showing an increase in this RBP in human DRG and peripheral nerves with a history of chronic neuropathic pain.

Overall, our study provides new, fundamental insight into human peripheral nerves and DPN. With increase in axonal loss, we observe a competitive interplay between regenerative and degenerative processes in DPN. On the one hand, there is an increase in immune cells and extracellular matrix as well as pro-inflammatory mediators that contribute to an inflammatory and degenerative phenotype. On the other hand, we detect mediators that can be targeted to activate regenerative pathways. Early targeting of genes and pathways involved in regenerative processes can open therapeutic avenues that directly treat DPN in peripheral nerves. Future studies using larger sample sizes and evaluating the functional effects of specific cells and mediators will be key to achieving this purpose. In addition, we identified the presence of specific mRNAs in the axons of human peripheral nerves, supporting our hypothesis that RNA transport occurs in human sensory nerves. It is likely that this is an important mechanism for the maintenance of axonal integrity and peripheral nerve homeostasis.

## MATERIALS AND METHODS

### Consent, tissue and patient data collection

All protocols were reviewed and approved by the UT Dallas (UTD) and UT Southwestern Medical Center (UTSWMC) Institutional Review Boards. Patients undergoing lower leg amputation at two major tertiary care hospitals (Clements University Hospital (CUH) at UTSWMC and Parkland Memorial Hospital (PMH)) in Dallas, Texas were recruited as part of the study. Informed consent for participation was obtained for each patient during study enrolment. Tibial and sural nerves were recovered from diabetic patients who have non-reconstructable soft tissue or bone loss, recalcitrant bone infection (osteomyelitis) and/or critical limb ischemia and had been advised to undergo lower extremity amputation. The tibial and sural nerves were harvested from the part of the leg which was amputated (distal specimen). The specimens were placed in sterile specimen cups, snap-frozen in liquid nitrogen and stored in a – 80C freezer. Control sural nerves were recovered from non-diabetic patients having lower leg amputation due to trauma or non-reconstructable deformity.

Details can be found in Suppl. File S1. Additional controls (peripheral nerves) were recovered from organ donors at Southwest Transplant Association within 4 hours of cross-clamp, frozen immediately on crushed dry ice, and stored in a -80°C freezer as previously described (104) (Suppl. Table S1). All human tissue procurement procedures from organ donors were also approved by the Institutional Review Boards at the University of Texas at Dallas.

### Sex as a biological variable

The demographics of our DPN population is 75% male (**Figure 1**; Suppl. File S1) limiting our analysis of sex-differences and interpretation of results. However, to account for potential differences due to sex, we included sex as a factor in our design formula for DESeq2 analysis.

### Tissue preparation

For Visium, Xenium, RNAscope and/or immunohistochemistry (IHC), tissues were gradually embedded in optimal cutting temperature (OCT) compound in a cryomold by adding small volumes of OCT over dry ice to avoid thawing. Nerves used for Visium and Xenium were cryosectioned onto Visium slides at 10µm. Tissues used for RNAscope and IHC were sectioned onto SuperFrost Plus charged slides at 20µm. Xenium protocol was performed at K2-Biolabs according to the manufacturer’s instructions.

### Peripheral nerve morphology

Frozen sural nerve biopsies received on dry ice were rapidly thawed for 1 minute at 37°C in a water bath, immediately fixed by immersion in 3% glutaraldehyde in 0.1 M phosphate buffer at room temperature for 12-15 hours, post-fixed in 1% osmium tetroxide for 2 hours, embedded in Epoxy resin, sectioned at 1 µm and stained with 1% Toluidine Blue (in 2% sodium borate in distilled water), as previously published (refs). Qualitative assessment of axonal density/loss was determined by a board-certified neuromuscular pathologist, based on clinical guidelines.

### RNAscope in situ hybridization

RNAscope in situ hybridization multiplex version 2 was performed as instructed by Advanced Cell Diagnostics (ACD) and as previously described (105). Optimal results were observed with protease digestion time of 10 seconds. Suppl. Tables S1 and S2 contain information on patients/donor tissues used and probes. All tissues were checked for RNA quality by using a positive control probe cocktail (ACD) which contains probes for high, medium and low-expressing mRNAs that are present in all cells (ubiquitin C > Peptidyl-prolyl cis-trans isomerase B > DNA-directed RNA polymerase II subunit RPB1). A negative control probe against the bacterial DapB gene (ACD) was used to reference non-specific/background label.

### Immunohistochemistry (IHC)

For dual RNAscope/IHC, after completion of RNAscope in situ hybridization, slides were incubated in blocking buffer (10% Normal Goat Serum, 0.3% Triton-X 100 in 0.1M PB) for 1 hour at room temperature while being shielded from light. Slides were placed in a light-protected humidity-controlled tray and incubated in primary antibody (Chicken Polyclonal Antibody to Peripherin, dilution 1:500, Encor Biotechnology, catalog number CPCA-Peri; Mouse Monoclonal Antibody to SOX10, dilution 1:40, Abcam, catalog number ab216020) in blocking buffer for 3 hours at room temperature or overnight at 4°C. Slides were washed with 0.1 M PB, then incubated in secondary antibody for 1 hour at room temperature. Slides were washed with 0.1 M PB, air-dried, and cover-slipped with Prolong Gold Antifade mounting medium. For regular IHC, slides were kept in the -20 C cryostat chamber for 15 minutes following completion of sectioning. The slides were then immediately fixed in ice-cold formalin (10%) for 1 minute followed by dehydration in 50% ethanol (1 minute), 70% ethanol (1 minute), and 100% ethanol (2 minutes) at room temperature. The slides were briefly air dried. A hydrophobic pen (ImmEdge PAP Pen; Vector Labs) was used to draw boundaries around each tissue section, and boundaries were allowed to air dry. Slides were incubated with blocking buffer (10% Normal Goat Serum, Atlanta Biologicals, Cat #S13150h, 0.3% Triton X-100 in 0.1 M PB) for 1 hour at room temperature. Sections were then incubated overnight with a primary antibody cocktail. Following primary antibody incubation, sections were washed with 0.1 M phosphate buffer and incubated with Alexa Fluor secondary antibodies (Fisher Scientific, dilutions 1:1000) for 1 hour at room temperature. Sections were washed in 0.1 M phosphate buffer. To remove lipofuscin signal, Trublack (1:20 in 70% ethanol; Biotium #23007) was pipetted to cover each section for 1 minute before being rinsed off. Finally, slides were air dried and cover slipped with Prolong Gold Antifade reagent (Fisher Scientific; P36930).

### Image acquisition

Sciatic and sural nerve sections were imaged on an Olympus FV3000 confocal microscope at 100X magnification. A minimum of 2 images were acquired for each nerve section. The area imaged was randomly chosen; however, we prioritized sections that did not have any sectioning artifact and sections that contained intact peripherin fiber staining. The acquisition parameters were set based on guidelines for the FV3000 provided by Olympus. Briefly, the gain was kept at the default setting 1, HV ≤ 600, offset was based on HI-LO settings, and laser power ≤ 5%.

### RNA-seq library preparation

#### Bulk RNA-sequencing

Following RNA purification, cDNA libraries were prepared with TruSeq Stranded Total RNA Library Prep with ribosomal RNA depletion for all samples according to the manufacturer’s instructions (Illumina). The quality of the extracted RNA and cDNA at each library preparation step was assessed with Qubit (Invitrogen) and High Sensitivity NGS fragment analysis kit on the Fragment Analyzer (Agilent Technologies). The amount of cDNA was standardized across samples and the libraries were sequenced on Illumina NextSeq500 sequencing machine with 75-bp single-end reads. mRNA library preparation and sequencing were done at the Genome Center in the University of Texas at Dallas Research Core Facilities.

#### Visium Spatial Gene Expression

Visium tissue optimization and spatial gene expression protocols were followed exactly as described by 10x Genomics (https://www.10xgenomics.com/) using Haematoxylin and Eosin as the counterstain. Optimal permeabilization time was obtained at 12 min incubation with permeabilization enzyme. Imaging was conducted on an Olympus vs120 slide scanner. Sural nerves from 5 patients were used. mRNA library preparation and sequencing were done at the Genome Center in the University of Texas at Dallas Research Core Facilities.

### RNA-seq – mapping raw counts and alignment of barcoded spots with imaged sections

#### Bulk

Sequenced reads were trimmed to avoid compositional bias and lower sequencing quality at either end and to ensure all quantified libraries were mapped with the same read length, and mapped to the GENCODE reference transcriptome (v27) in a strand-aware and splicing-aware fashion using the STAR alignment tool (106). Stringtie (107) was used to generate relative abundances in Transcripts per Million (TPM), and non-mitochondrial coding gene abundances were extracted and renormalized to a million to generate coding TPMs for each sample.

#### Visium

The output data of each sequencing run (Illumina BCL files) was processed using the Space Ranger (v1.1) pipelines provided by 10x Genomics. Samples were demultiplexed into FASTQ files using Space Ranger’s mkfastq pipeline. Space Ranger’s count pipeline was used to align FASTQ files with brightfield microscope images previously acquired, detect barcode/UMI counting, and map reads to the human reference transcriptome (Gencode v27 and GRCh38.p10) (108). This pipeline generates, for each sample, feature-barcode matrices that contain raw counts and places barcoded spots in spatial context on the slide image (cloupe files). Gene expression with spatial context can, then, be visualized by loading cloupe files onto Loupe Browser (v5, 10x Genomics).

### Somascan Assay

Tissue lysates were prepared from fresh-frozen DRGs and peripheral nerves. The tissues were placed in T-PER Tissue Protein Extraction Reagent (Thermo Scientific, Cat # 78510) with additional 1X Halt Protease Inhibitor Cocktail (Thermo Scientific, Cat # 87786) and homogenized using Precellys Soft Tissue Homogenizing beads (Bertin Corp, Cat # P000933-LYSK0-A.0). Samples were centrifuged at 14,000 x g for 15 minutes in the cold room. The resulting supernatant was quantified Micro BCA™ Protein Assay Kit (Thermo Scientific, Cat# 23235) and normalized accordingly. Proteins were profiled using the SOMAScan platform. 7000 analytes were measured on the the SOMAScan assay. Quality controls were performed by SomaLogic to correct for technical variabilities within-run and between-run for each sample.

### Statistics: Differential expression analysis

We performed differential expression analysis using the “DESeq” function (this function performs differential expression analysis based on the negative binominal distribution and Wald statistics). Nominal P values were corrected for multiple testing using the Benjamini-Hochberg (BH) method (85). In addition, we performed shrinkage of the log2 FC (LFC) estimates to generate more accurate LFC. We used the adaptive shrinkage estimator from the “ashr” R package (86) and set the contrast to ‘sural’ vs ‘tibial’ or ‘moderate’ versus ‘severe’ as the groups we wanted to compare. Genes were considered to be DE if FC > 1.5 and adjusted P < 0.05. Statistical hypothesis testing results for all tests can be found in **Suppl. Files S3** and **S5**. For each gene tested, we report baseMean (mean of normalized counts), LFC, lfcSE (standard error of the LFC estimate), P value (Wald test P value), and Padj (BH-adjusted P values).

### Statistics: Analysis of visium spatial transcriptomics data

We used Python (v3.8 with Anaconda distribution), R (v4.3.1), and Seurat (v4.3.0) (109) for data analysis. We used Seurat’s spatially-resolved RNA-seq data analysis workflow and used H5 files (output of the 10x spaceranger pipeline) as input for the spatial analysis. This analysis follows Seurat’s standard graph-based clustering with the addition of tools for the integration and visualization of spatial and molecular information. The workflow consists of data normalization (using sctransform(110)), dimensional reduction and clustering, and identification of spatially-variable features that we used to label the clusters. The proportion of enriched cell groups was computed used the package DittoSeq (111). For spot deconvolution, we used STdeconvolve (46), a reference-free deconvolution tool. Cell types were identified based the top markers in each topic.

### Statistics: Cell-cell interactions

We used the standard Cellchat (45) workflow to identify cell-cell interactions between cell types enriched in moderate and severe axonal loss samples. Separate CellChat objects for each sample were generated according to the instructions in the CellChat manual, using the createCellChat() function. This involved incorporating the normalized, scaled RNA data assay and the annotation labels from each corresponding Seurat object. Following the CellChat methodology, key steps included identifying the genes and interactions that were overexpressed in each cell group using identifyOverExpressedGenes() and identifyOverExpressedInteractions(). The computation of communication probability was carried out using computeCommunProb(), adhering to default settings. This function employs the trimean method, under the assumption that the mean expression level of a gene is considered zero if expressed in less than 25% of cells, thus resulting in a reduced number of interactions but enhancing statistical reliability. The communication network underwent refinement via filterCommunication(), which by default excludes cell groups comprising fewer than 10 cells.

Assessment of communication at the pathway level was conducted with computeCommunProbPathway(). Visualization of the interactions utilized CellChat’s suite of plotting tools, which include chord plots, circos plots, and heatmaps.

## Study approval

This work was approved by the University of Texas at Dallas Institutional Review Board (MR-19-022 – tibial and sural nerves from amputation surgeries and MR-15-237 – DRG and peripheral nerves from organ donors).

## Data availability

Public access to processed data will be available at sensoryomics.com.

## AUTHOR CONTRIBUTIONS

DT-F, TL, DW and TJP designed the study. DT-F performed RNA extractions for bulk-RNA sequencing. DT-F, SS and IS performed visium experiments. DT-F, SS and BQS performed RNAscope and IHC experiments. JMM performed SomaScan assay and analysis. DT-F analysed sequencing data. MK, NI and KM assisted in the analysis of bulk RNA sequencing. EEU performed sural nerve morphology and qualitatively determined axonal density. TL, DW and GT collected tibial and sural nerve samples. GT coordinated clinical aspects of the study. DW and TJP supervised the study. DT-F, BQS and TJP wrote the manuscript. All authors read and edited the paper.

## ACKNOWLEDGEMENTS

The authors thank the Genome Center at The University of Texas at Dallas for the services to support our research. We thank the patients for participating in this study. We also thank Erin Vines, Anna Cervantes, Geoffrey Funk and Peter Horton at Southwest Transplant Alliance and the organ donors and their families for their generous gift. We extend our thanks also to all members of the Price lab for participating in organ donor recoveries. This work was supported by NIH grants NS111929 and NS065926 to TJP.

