## Supplementary material for "Deciphering the molecular landscape of human peripheral nerves: implications for diabetic peripheral neuropathy": Suppl. Figures and Tables

#### SUPPLEMENTARY MATERIALS

##### Supplementary figures:

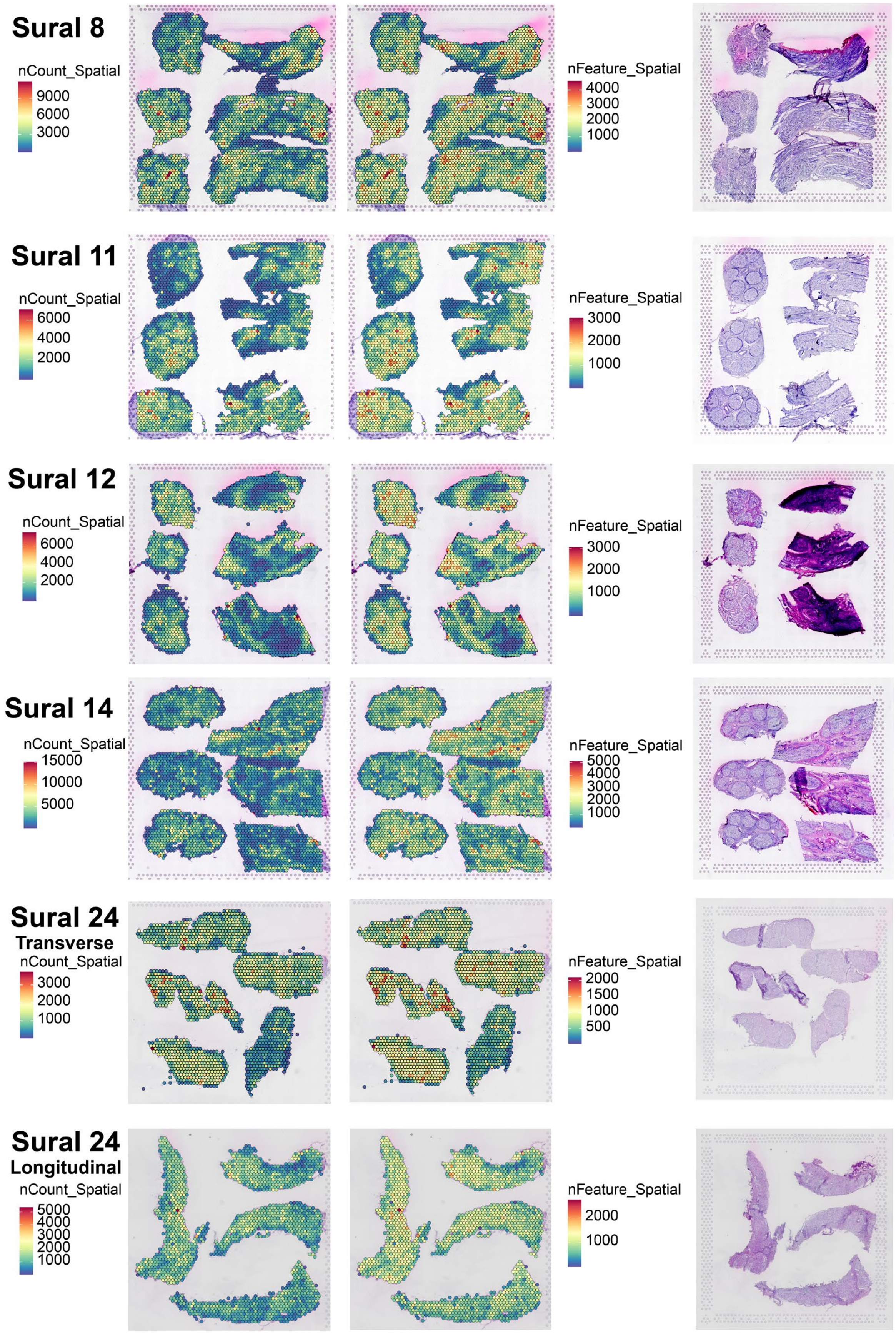

Sural 8
nCount_Spatial
9000
6000
3000
nFeature_Spatial
4000
3000
2000
1000
Sural 11
nCount_Spatial
6000
4000
2000
nFeature_Spatial
3000
2000
1000
Sural 12
nCount_Spatial
6000
4000
2000
nFeature_Spatial
3000
2000
1000
Sural 14
nCount_Spatial
15000
10000
5000
nFeature_Spatial
5000
4000
3000
2000
1000
Sural 24
Transverse
nCount_Spatial
3000
2000
1000
nFeature_Spatial
2000
1500
1000
500
Sural 24
Longitudinal
nCount_Spatial
5000
4000
3000
2000
1000
nFeature_Spatial
2000
1000

**Suppl. Figure 1. Quality control of sural nerves.** Plots display number of reads (nCount_Spatial) and number of genes (nFeature_Spatial) next to respective H&E image for each sural nerve.

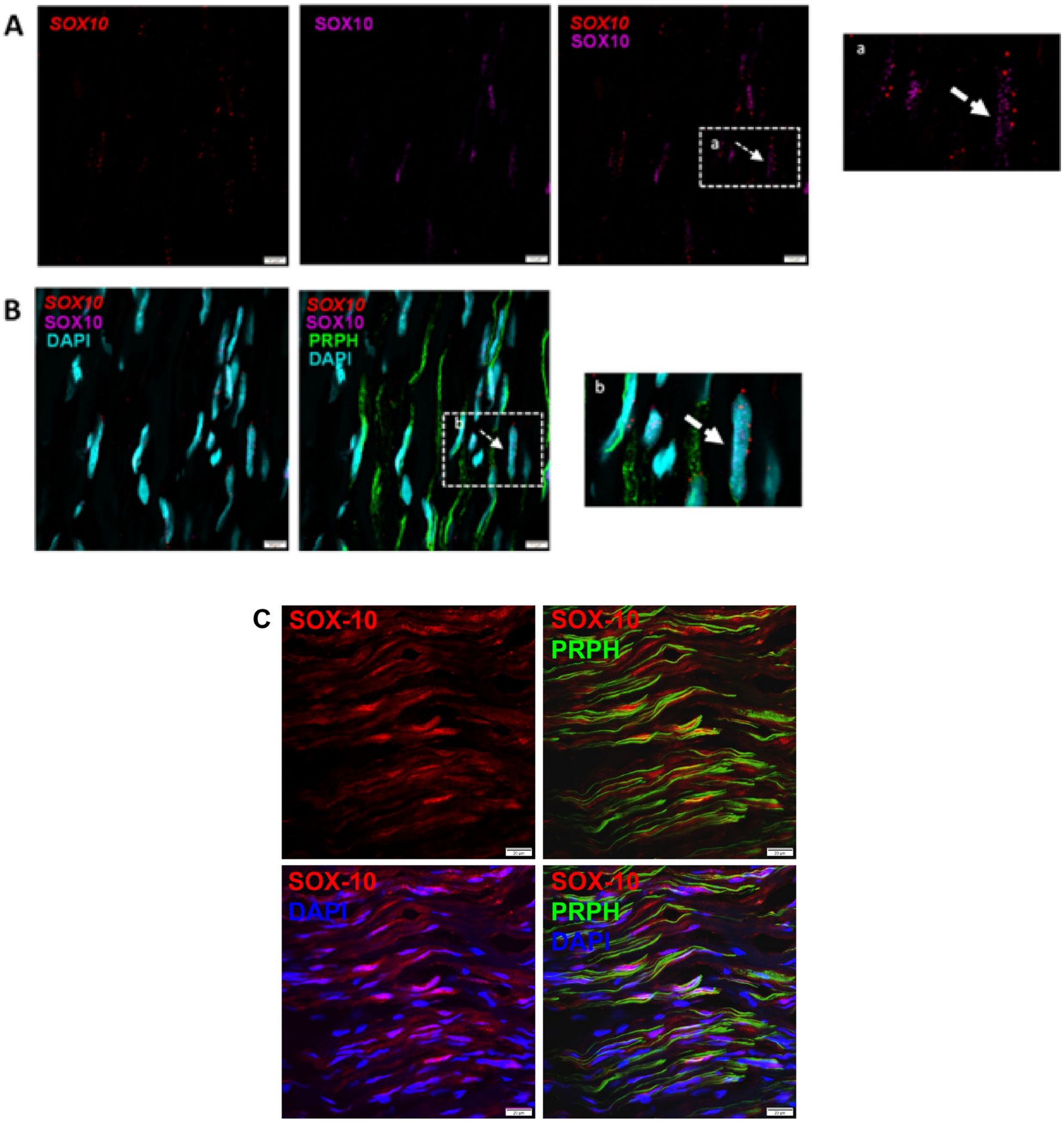


**Suppl. Figure 2. SOX10+ Schwann cells are present in human peripheral nerves**. **A**) *SOX10* mRNA puncta (red) colocalize with SOX10 protein (magenta). **B**) *SOX10* mRNA puncta (red) colocalize with DAPI (cyan). Peripherin (PRPH, green) was used to label nerve fibers. Insets show zoomed-in images. Scale bars=10 µm. C) Expression of SOX10 protein using a second antibody. Scale bars=10 µm.

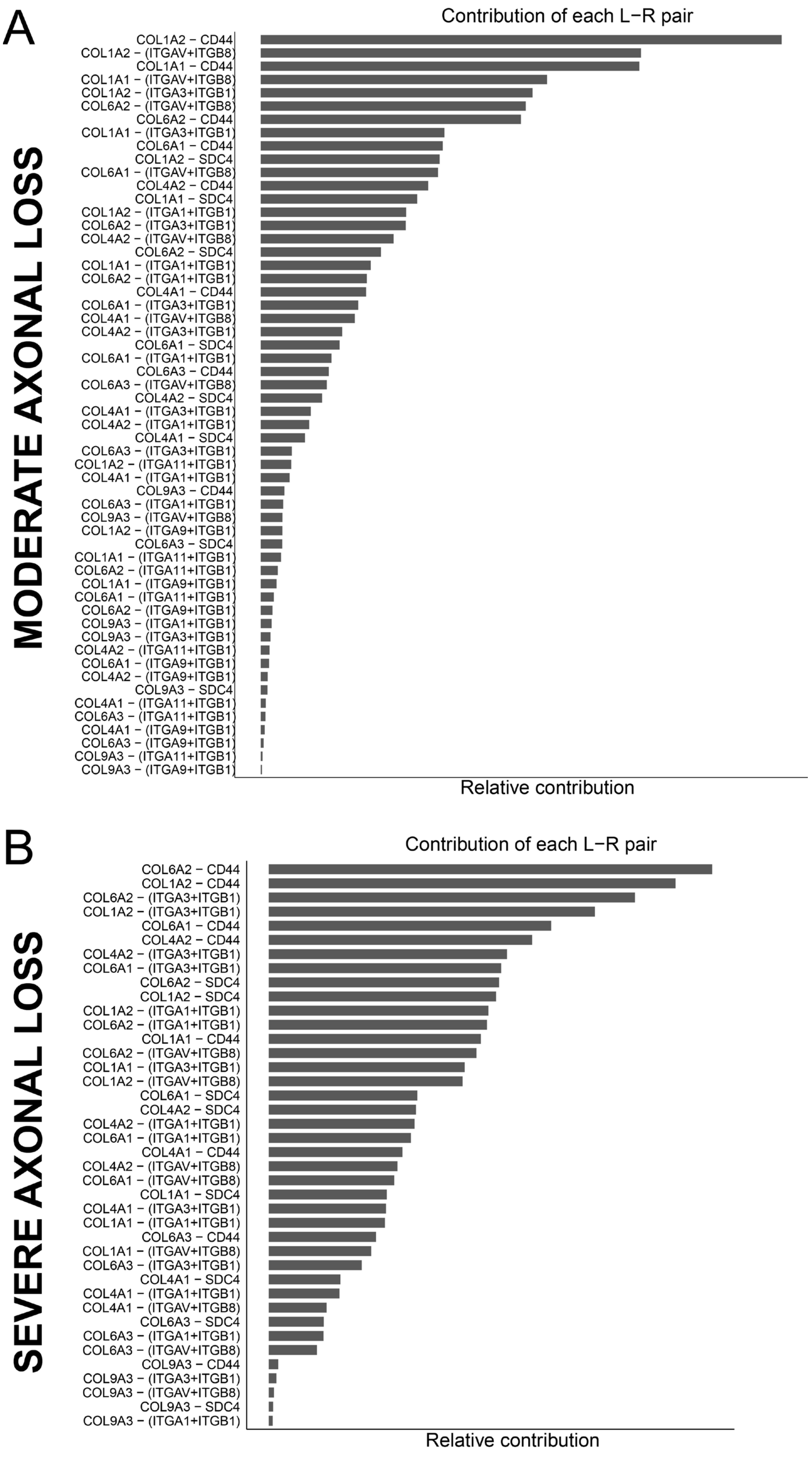
A
MODERATE AXONAL LOSS
Contribution of each L−R pair
COL1A2 − CD44
COL1A2 − (ITGAV+ITGB8)
COL1A1 − CD44
COL1A1 − (ITGAV+ITGB8)
COL1A2 − (ITGA3+ITGB1)
COL6A2 − (ITGAV+ITGB8)
COL6A2 − CD44
COL1A1 − (ITGA3+ITGB1)
COL6A1 − CD44
COL1A2 − SDC4
COL6A1 − (ITGAV+ITGB8)
COL4A2 − CD44
COL1A1 − SDC4
COL1A2 − (ITGA1+ITGB1)
COL6A2 − (ITGA3+ITGB1)
COL4A2 − (ITGAV+ITGB8)
COL6A2 − SDC4
COL1A1 − (ITGA1+ITGB1)
COL6A2 − (ITGA1+ITGB1)
COL4A1 − CD44
COL6A1 − (ITGA3+ITGB1)
COL4A1 − (ITGAV+ITGB8)
COL4A2 − (ITGA3+ITGB1)
COL6A1 − SDC4
COL6A1 − (ITGA1+ITGB1)
COL6A3 − CD44
COL6A3 − (ITGAV+ITGB8)
COL4A2 − SDC4
COL4A1 − (ITGA3+ITGB1)
COL4A2 − (ITGA1+ITGB1)
COL4A1 − SDC4
COL6A3 − (ITGA3+ITGB1)
COL1A2 − (ITGA11+ITGB1)
COL4A1 − (ITGA1+ITGB1)
COL9A3 − CD44
COL6A3 − (ITGA1+ITGB1)
COL9A3 − (ITGAV+ITGB8)
COL1A2 − (ITGA9+ITGB1)
COL6A3 − SDC4
COL1A1 − (ITGA11+ITGB1)
COL6A2 − (ITGA11+ITGB1)
COL1A1 − (ITGA9+ITGB1)
COL6A1 − (ITGA11+ITGB1)
COL6A2 − (ITGA9+ITGB1)
COL9A3 − (ITGA1+ITGB1)
COL9A3 − (ITGA3+ITGB1)
COL4A2 − (ITGA11+ITGB1)
COL6A1 − (ITGA9+ITGB1)
COL4A2 − (ITGA9+ITGB1)
COL9A3 − SDC4
COL4A1 − (ITGA11+ITGB1)
COL6A3 − (ITGA11+ITGB1)
COL4A1 − (ITGA9+ITGB1)
COL6A3 − (ITGA9+ITGB1)
COL9A3 − (ITGA11+ITGB1)
COL9A3 − (ITGA9+ITGB1)
Relative contribution
B
SEVERE AXONAL LOSS
Contribution of each L−R pair
COL6A2 − CD44
COL1A2 − CD44
COL6A2 − (ITGA3+ITGB1)
COL1A2 − (ITGA3+ITGB1)
COL6A1 − CD44
COL4A2 − CD44
COL4A2 − (ITGA3+ITGB1)
COL6A1 − (ITGA3+ITGB1)
COL6A2 − SDC4
COL1A2 − SDC4
COL1A2 − (ITGA1+ITGB1)
COL6A2 − (ITGA1+ITGB1)
COL1A1 − CD44
COL6A2 − (ITGAV+ITGB8)
COL1A1 − (ITGA3+ITGB1)
COL1A2 − (ITGAV+ITGB8)
COL6A1 − SDC4
COL4A2 − SDC4
COL4A2 − (ITGA1+ITGB1)
COL6A1 − (ITGA1+ITGB1)
COL4A1 − CD44
COL4A2 − (ITGAV+ITGB8)
COL6A1 − (ITGAV+ITGB8)
COL1A1 − SDC4
COL4A1 − (ITGA3+ITGB1)
COL1A1 − (ITGA1+ITGB1)
COL6A3 − CD44
COL1A1 − (ITGAV+ITGB8)
COL6A3 − (ITGA3+ITGB1)
COL4A1 − SDC4
COL4A1 − (ITGA1+ITGB1)
COL4A1 − (ITGAV+ITGB8)
COL6A3 − SDC4
COL6A3 − (ITGA1+ITGB1)
COL6A3 − (ITGAV+ITGB8)
COL9A3 − CD44
COL9A3 − (ITGA3+ITGB1)
COL9A3 − (ITGAV+ITGB8)
COL9A3 − SDC4
COL9A3 − (ITGA1+ITGB1)
Relative contribution

**Suppl. Figure 3. Ligands and receptors within the Collagen signaling pathway.**
Contribution of each ligand-receptor pair in moderate **(A)** and severe **(B)** axonal loss nerves.

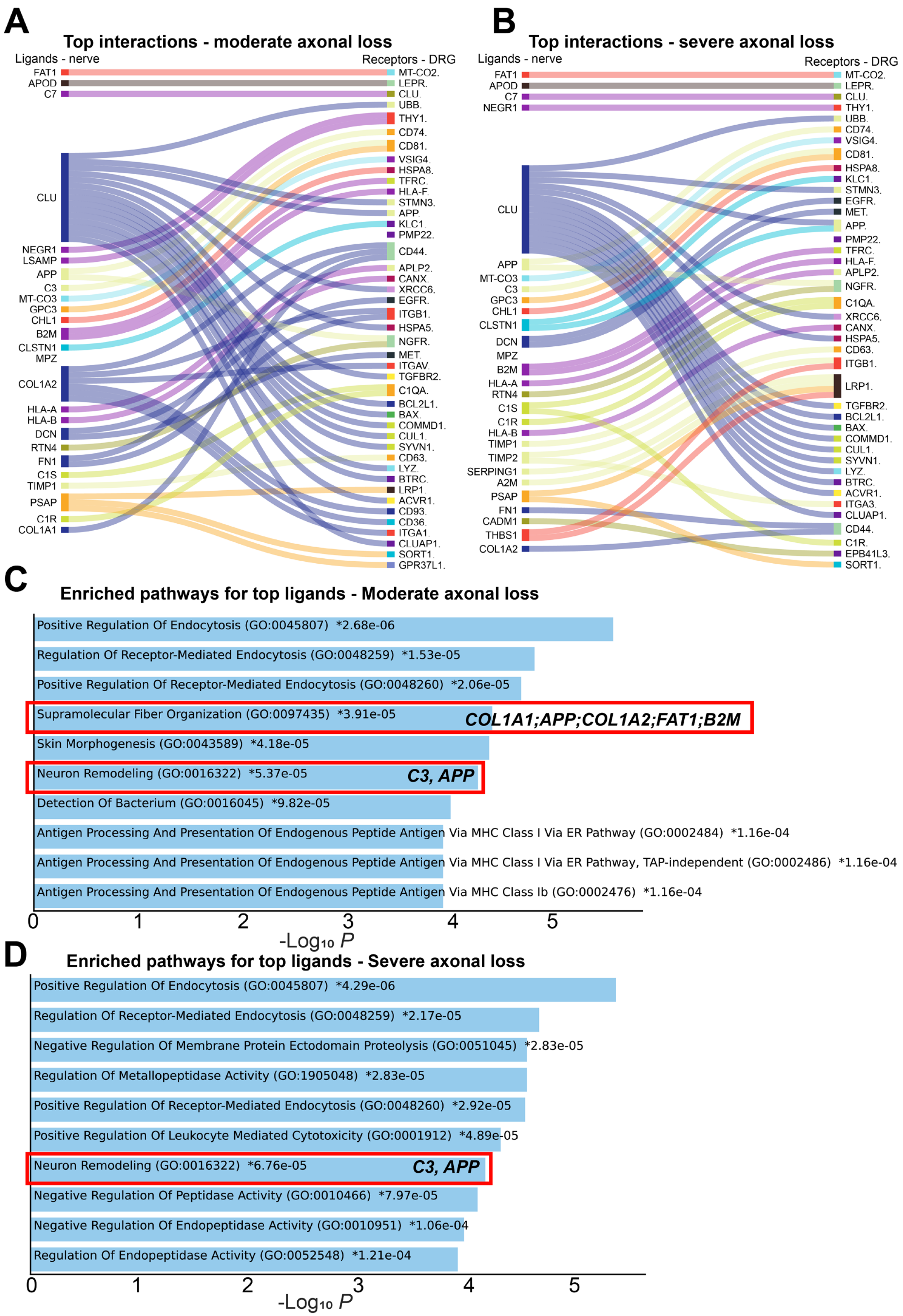
A
Top interactions - moderate axonal loss
Ligands - nerve
Receptors - DRG
FAT1
APOD
C7
CLU
NEGR1
LSAMP
APP
C3
MT-CO3
GPC3
CHL1
B2M
CLSTN1
MPZ
COL1A2
HLA-A
HLA-B
DCN
RTN4
FN1
C1S
TIMP1
PSAP
C1R
COL1A1
MT-CO2.
LEPR.
CLU.
UBB.
THY1.
CD74.
CD81.
VSIG4.
HSPA8.
TFRC.
HLA-F.
STMN3.
APP.
KLC1.
PMP22.
CD44.
APLP2.
CANX.
XRCC6.
EGFR.
ITGB1.
HSPA5.
NGFR.
MET.
ITGAV.
TGFBR2.
C1QA.
BCL2L1.
BAX.
COMMD1.
CUL1.
SYVN1.
CD63.
LYZ.
BTRC.
LRP1.
ACVR1.
CD93.
CD36.
ITGA1.
CLUAP1.
SORT1.
GPR37L1.
B
Top interactions - severe axonal loss
Ligands - nerve
Receptors - DRG
FAT1
APOD
C7
NEGR1
CLU
APP
MT-CO3
C3
GPC3
CHL1
CLSTN1
DCN
MPZ
B2M
HLA-A
RTN4
C1S
C1R
HLA-B
TIMP1
TIMP2
SERPING1
A2M
PSAP
FN1
CADM1
THBS1
COL1A2
MT-CO2.
LEPR.
CLU.
THY1.
UBB.
CD74.
VSIG4.
CD81.
HSPA8.
KLC1.
STMN3.
EGFR.
MET.
APP.
PMP22.
TFRC.
HLA-F.
APLP2.
NGFR.
C1QA.
XRCC6.
CANX.
HSPA5.
CD63.
ITGB1.
LRP1.
TGFBR2.
BCL2L1.
BAX.
COMMD1.
CUL1.
SYVN1.
LYZ.
BTRC.
ACVR1.
ITGA3.
CLUAP1.
CD44.
C1R.
EPB41L3.
SORT1.
C
Enriched pathways for top ligands - Moderate axonal loss
Positive Regulation Of Endocytosis (GO:0045807) *2.68e-06
Regulation Of Receptor-Mediated Endocytosis (GO:0048259) *1.53e-05
Positive Regulation Of Receptor-Mediated Endocytosis (GO:0048260) *2.06e-05
Supramolecular Fiber Organization (GO:0097435) *3.91e-05
COL1A1;APP;COL1A2;FAT1;B2M
Skin Morphogenesis (GO:0043589) *4.18e-05
Neuron Remodeling (GO:0016322) *5.37e-05
C3, APP
Detection Of Bacterium (GO:0016045) *9.82e-05
Antigen Processing And Presentation Of Endogenous Peptide Antigen Via MHC Class I Via ER Pathway (GO:0002484) *1.16e-04
Antigen Processing And Presentation Of Endogenous Peptide Antigen Via MHC Class I Via ER Pathway, TAP-independent (GO:0002486) *1.16e-04
Antigen Processing And Presentation Of Endogenous Peptide Antigen Via MHC Class Ib (GO:0002476) *1.16e-04
0
1
2
3
4
5
-Log10 P
D
Enriched pathways for top ligands - Severe axonal loss
Positive Regulation Of Endocytosis (GO:0045807) *4.29e-06
Regulation Of Receptor-Mediated Endocytosis (GO:0048259) *2.17e-05
Negative Regulation Of Membrane Protein Ectodomain Proteolysis (GO:0051045) *2.83e-05
Regulation Of Metallopeptidase Activity (GO:1905048) *2.83e-05
Positive Regulation Of Receptor-Mediated Endocytosis (GO:0048260) *2.92e-05
Positive Regulation Of Leukocyte Mediated Cytotoxicity (GO:0001912) *4.89e-05
Neuron Remodeling (GO:0016322) *6.76e-05
C3, APP
Negative Regulation Of Peptidase Activity (GO:0010466) *7.97e-05
Negative Regulation Of Endopeptidase Activity (GO:0010951) *1.06e-04
Regulation Of Endopeptidase Activity (GO:0052548) *1.21e-04
0
1
2
3
4
5
-Log10 P

**Suppl. Figure 4. Interactome analysis between ligands enriched in the nerve and DRG receptors. A)** Interactome plot showing interactions between top ligands expressed in a sample with moderate axonal loss. **B)** Interactome plot showing interactions between top ligands expressed in a sample with severe axonal loss. **C, D)** Enriched pathways for top ligands in moderate **(C)** and severe **(D)** axonal loss.

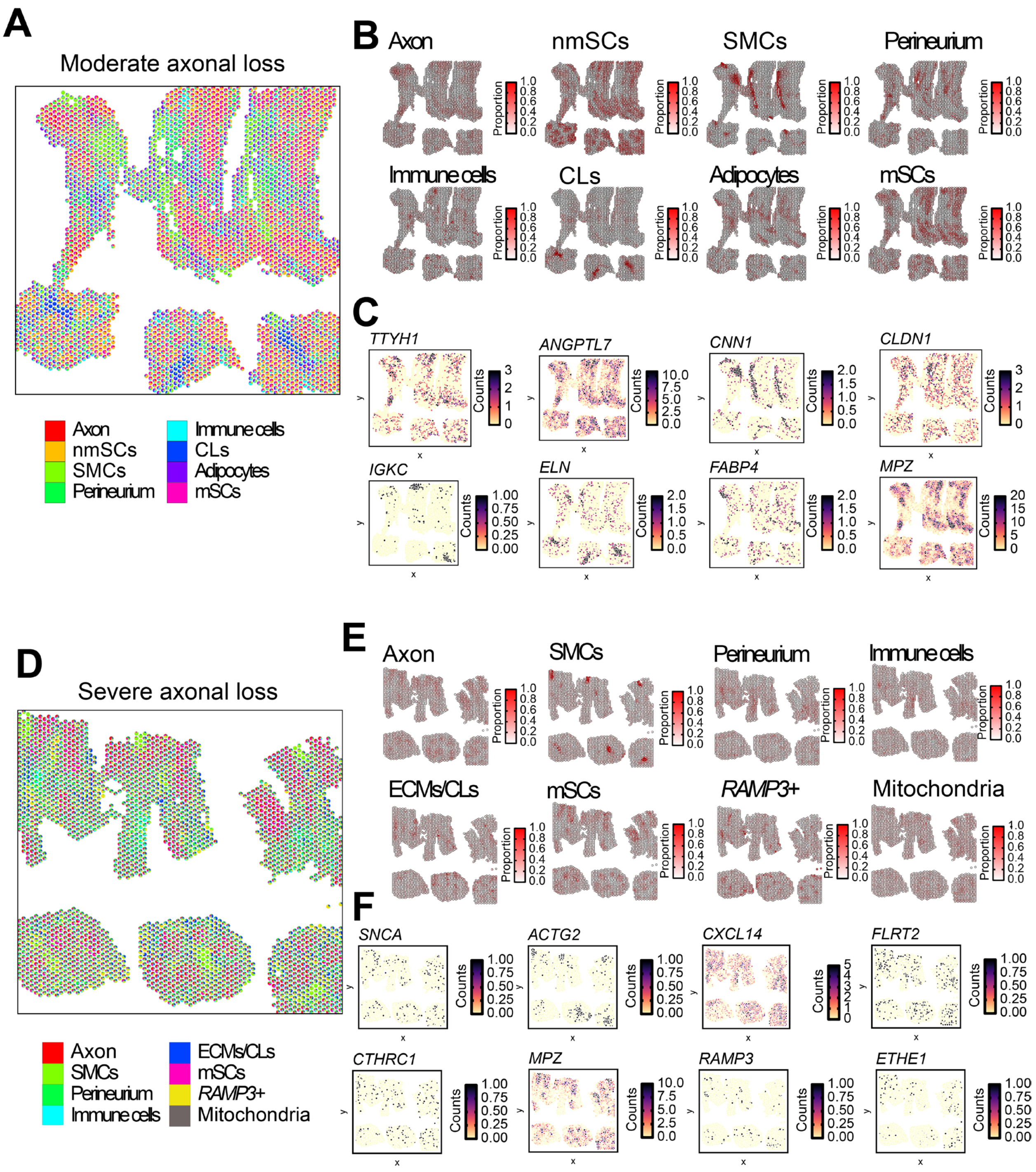


**Suppl. Figure 5. Cell type deconvolution of moderate and severe axonal loss sural nerves**. Deconvolution of cells in a sample with moderate axonal loss **(A)** and in a sample with severe axonal loss **(D)**. Proportion of cells identified in the sample with moderate axonal loss **(B)** and in the sample with severe axonal loss **(E)**. Markers for each cell type in sample with moderate axonal loss **(C)** and with severe axonal loss **(F)**. SMCs=Smooth muscle cells; ECM=Extracellular matrix; CLs= Connective layers; nmSCs= non-myelinating Schwann cells; mSCs= myelinating Schwann cells.

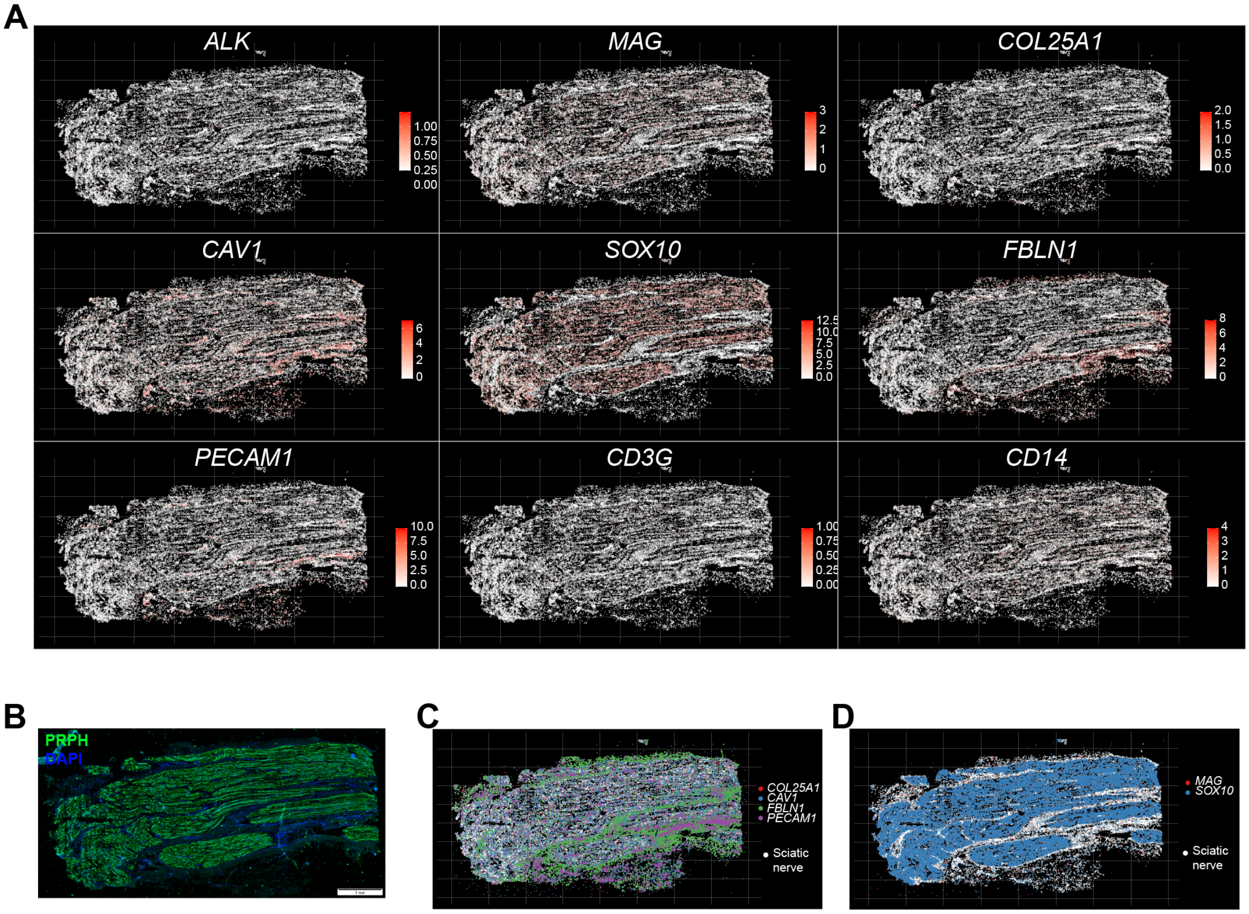


**Suppl. Figure 6. Visualization of specific genes in human peripheral nerves using Xenium.** A) Spatial localization of genes differentially expressed between moderate and severe axonal loss: ALK receptor tyrosine kinase (*ALK*), myelin associated glycoprotein (*MAG*), calveolin-1 (*CAV1*) and Collagen Type XXV Alpha 1 Chain (*COL25A1*) and cell type markers (SOX10-Schwann cells, PECAM1-endothelial cells, FBLN1-fibroblasts, CD3G-T cells, CD14-macrophages). B) Immunohistochemistry staining showing nerve fibers labeled with PRPH (green) and DAPI (blue). C) Co-localization of *CAV1* and *COL25A1* with fibroblast and endothelial cell markers. D) Co-localization of *MAG* with Schwann cell marker.

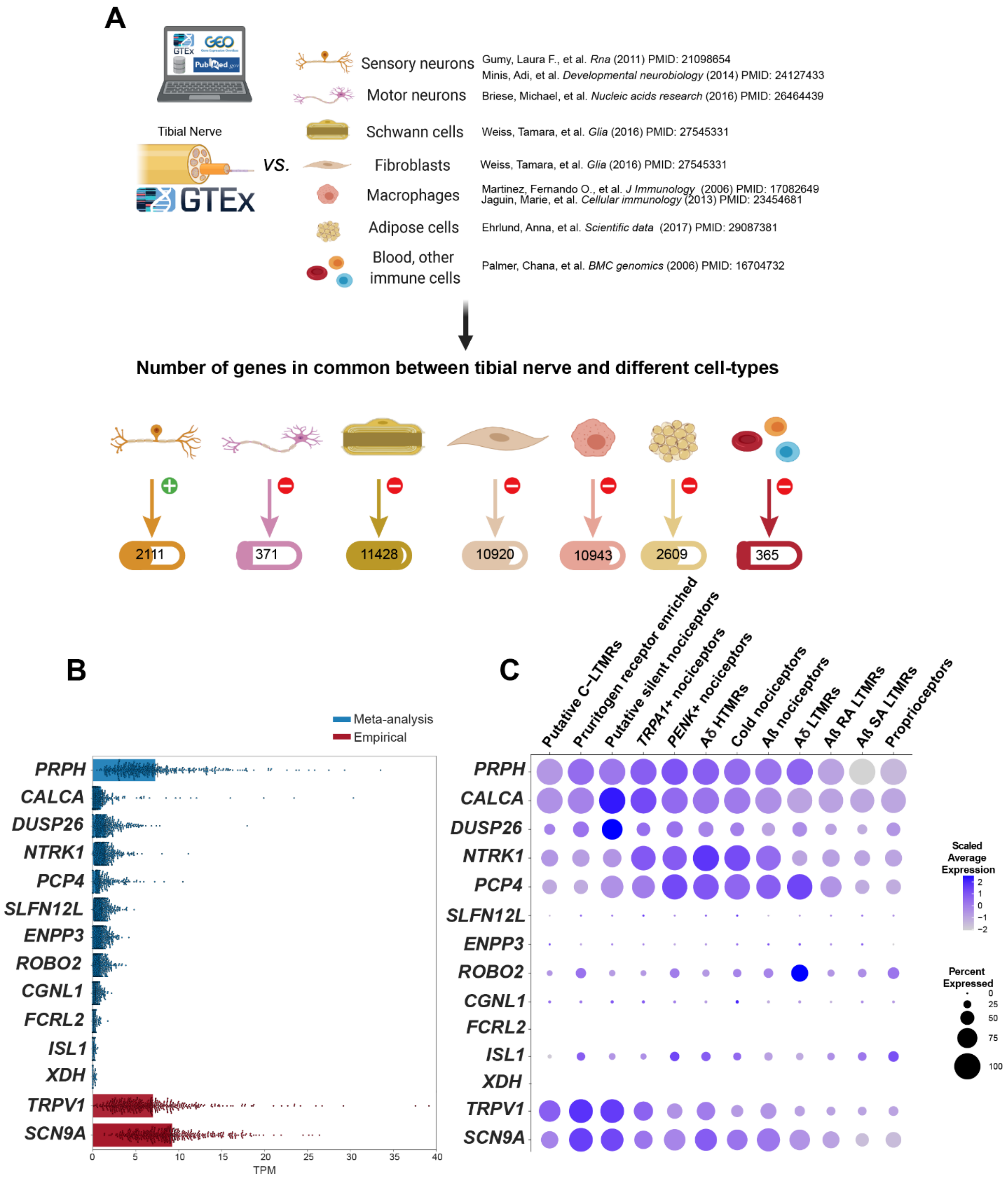


**Suppl. Figure 7. Meta-analysis of putative axonal mRNAs. A)** Diagram of publicly available studies included in meta-analysis. **B)** List of genes that were present only in sensory axons and that are likely axonal mRNAs (blue). In red are genes that are not exclusively axonal. **C)** Dot plot showing how putative axonal genes are expressed in human dorsal root ganglia (DRG) neurons. The size of the dot represents the percentage of barcodes within a cluster, and the color corresponds to the average expression (scaled data) across all barcodes within a cluster for each gene shown.

### Sciatic Nerve

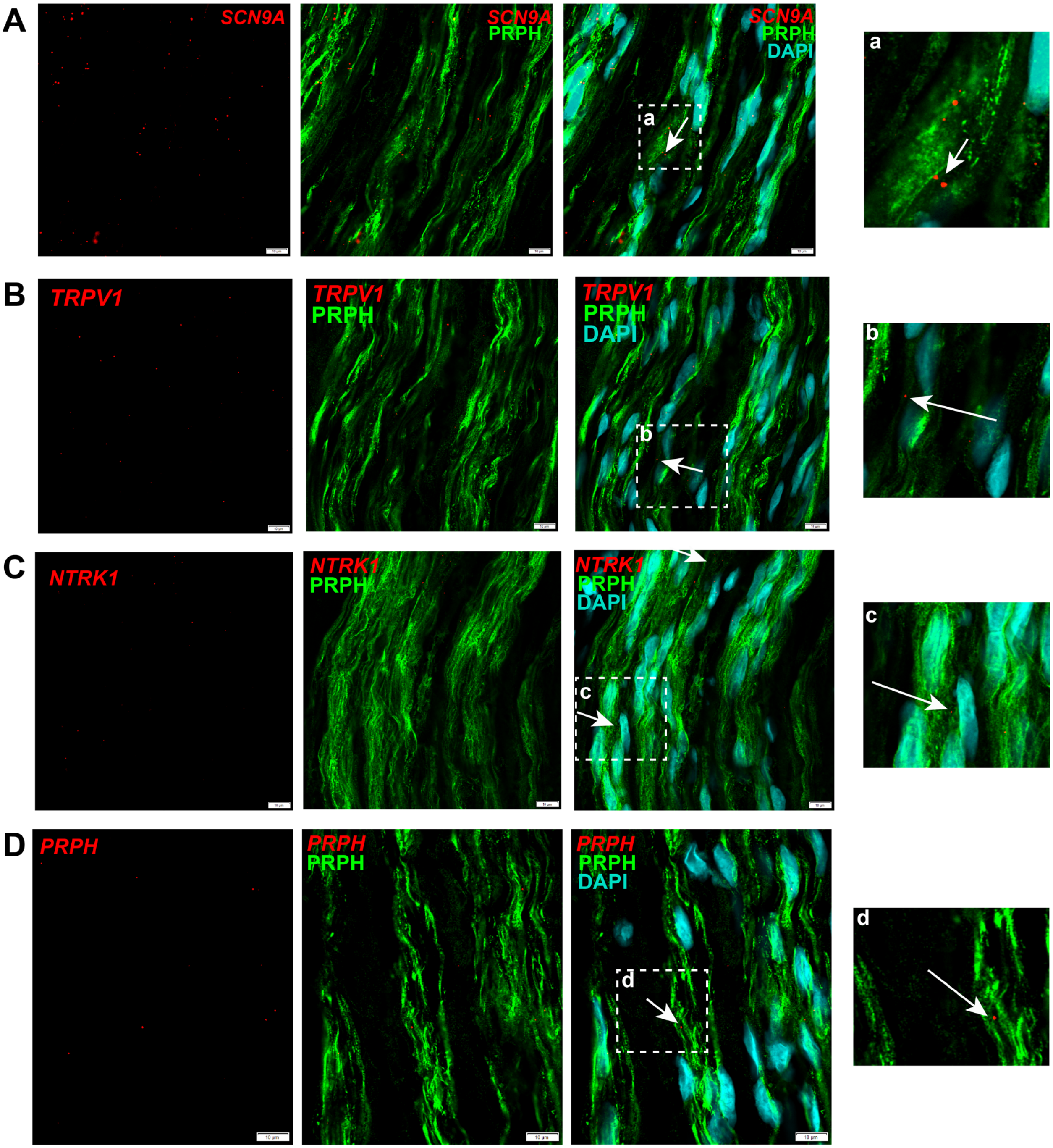


**Suppl. Figure 8. Characterization of Sciatic Nerve mRNAs with RNAscope and IHC**. mRNA puncta of *SCN9A* **(A)**, *TRPV1* **(B)**, *NTRK1* **(C)**, *PRPH* **(D)** in red are colocalized with peripherin (PRPH, green), which labels nerve fibers. Arrows point to areas where mRNA puncta do not overlap with DAPI (cyan), suggesting that it is axonal specific staining. Insets show zoomed in images. Scale bars=10 µm.

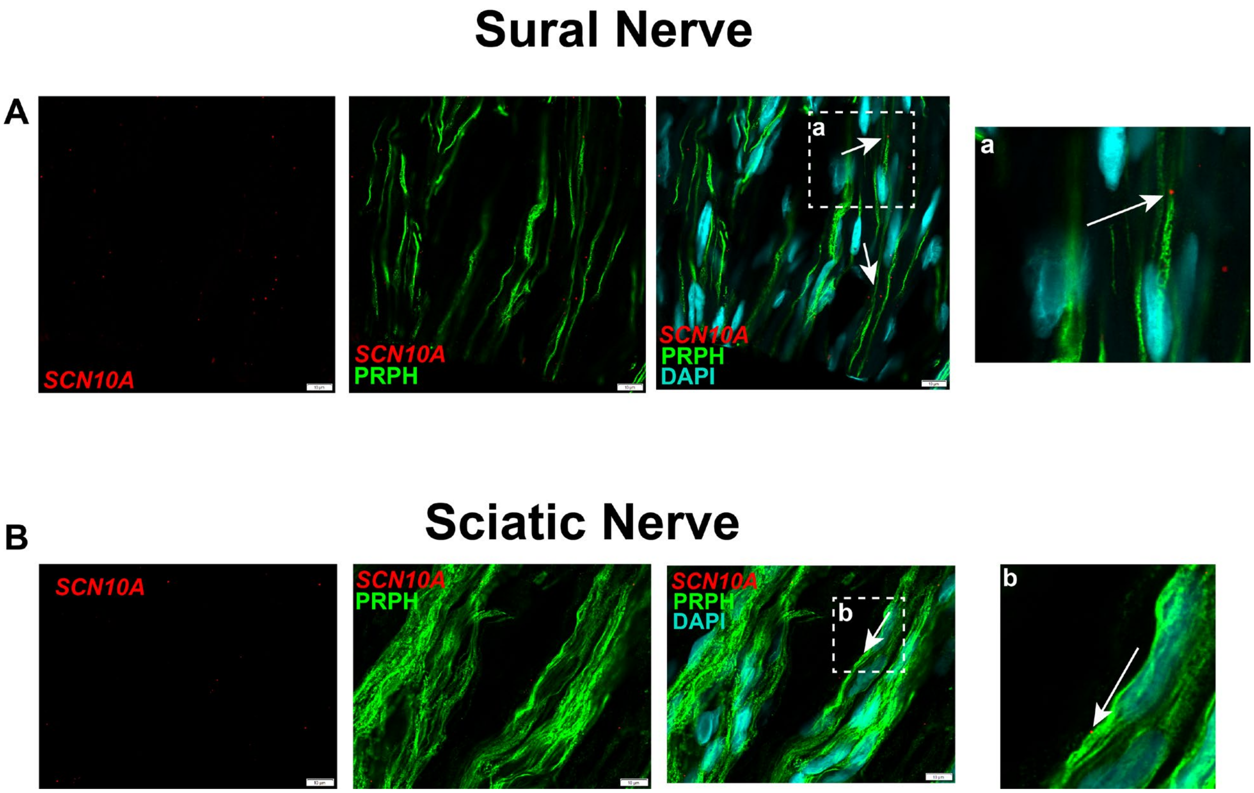


**Suppl. Figure 9. Presence of SCN10A punta in sural and sciatic nerve.** mRNA puncta of *SCN10A* in sural nerve **(A)** and sciatic nerve **(B)** in red are colocalized with peripherin (PRPH, green), which labels nerve fibers. Arrows point to areas where mRNA puncta do not overlap with DAPI (cyan), suggesting that it is axonal specific staining. Insets show zoomed in images. Scale bars=10 μm.

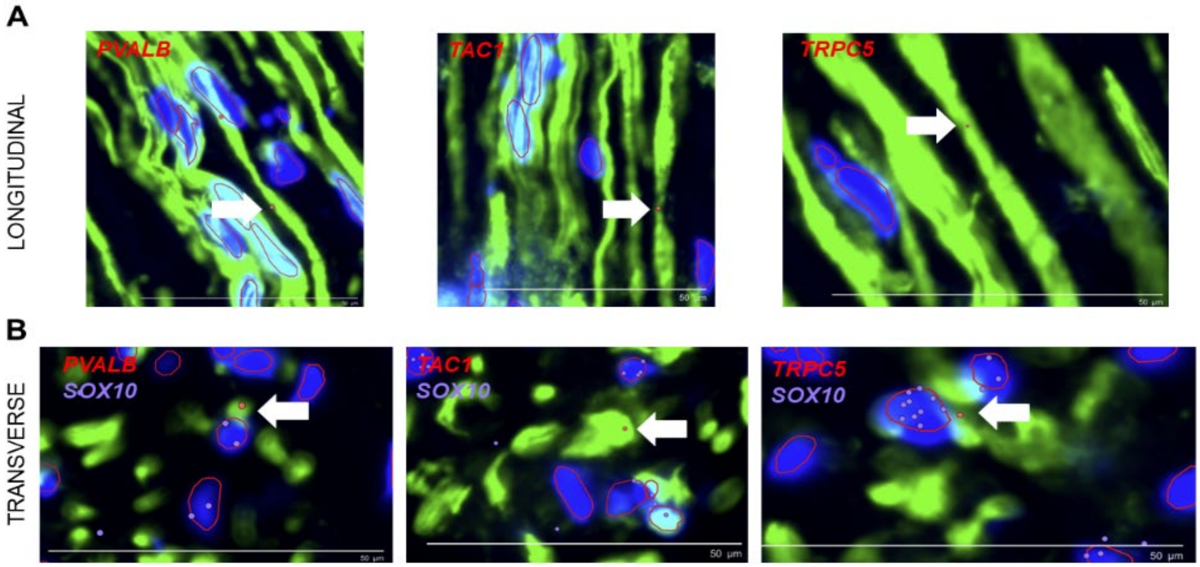


**Suppl. Figure 10. Axonal mRNAs are detected using Xenium in situ**. We used the off-the-shelf Xenium Brain panel and detect the presence of neuronal markers parvalbumin (*PVALB*), tachykinin precursor 1 (*TAC1*) and transient receptor potential cation channel subfamily c member 5 (*TRPC5*) in human peripheral axons (labeled by peripherin in green, PRPH) in longitudinal **(A)** and transverse **(B)** sections. SRY-Box Transcription Factor 10 (*SOX10*) is a marker of Schwann cells and its expression in DAPI+ cells surrounding the nerve fibers was used to assist in the validation of our cell segmentation. Scale bar= 50 μm.

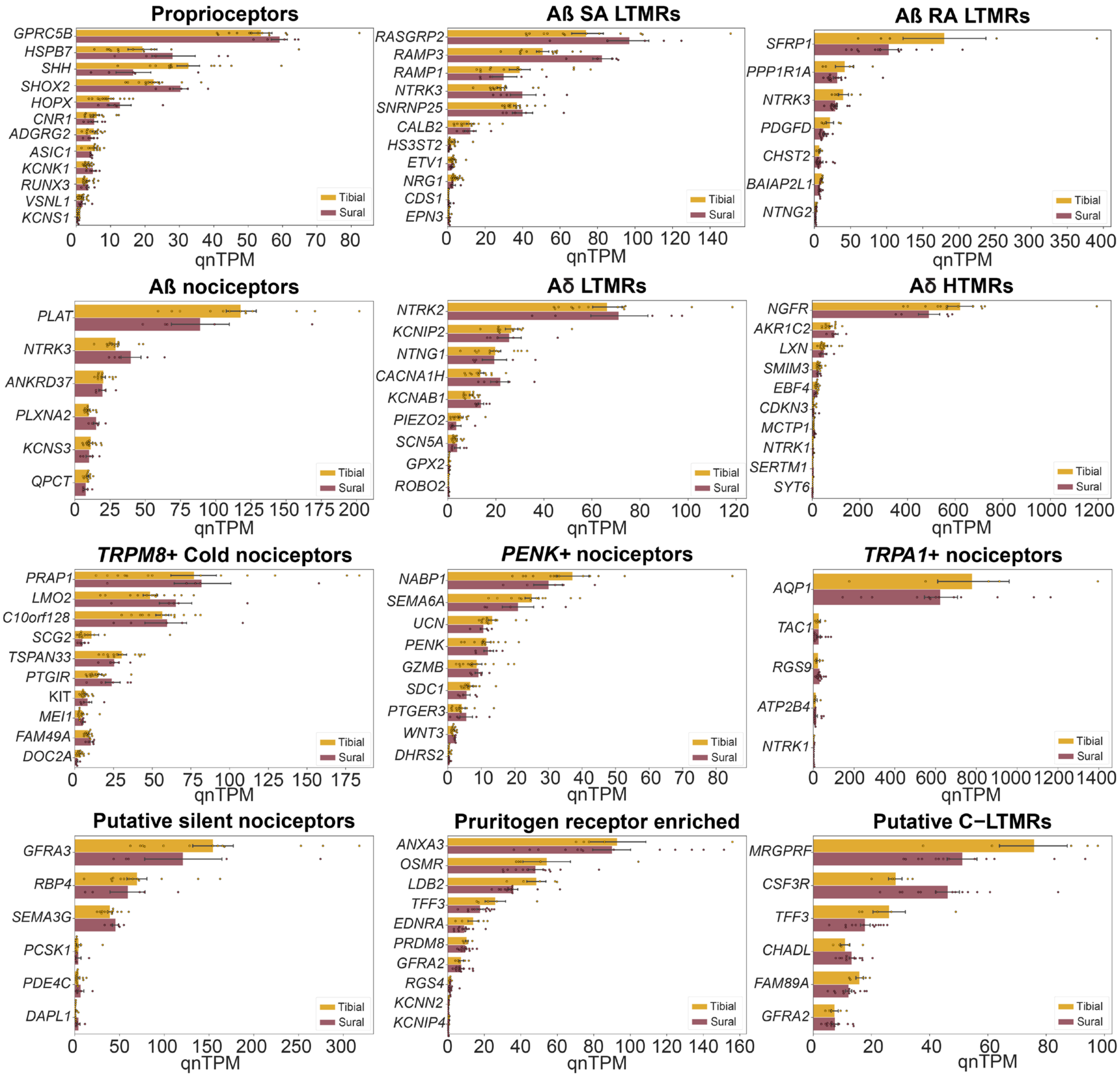


**Suppl. Figure 11. Markers of different neuronal subtypes can be identified in human sural and tibial nerves.**

**Suppl. Tables:**

**Table S1:** Tissue Donor/Patient Information for RNAscope/IHC.

| Donor/Patient | Tissue | Sex | Age | Cause of Death | Diabetes |
|---|---|---|---|---|---|
| D1 | Peripheral nerve | M | 37 | Anoxia/overdose | No |
| D2 | Peripheral nerve | F | 38 | Anoxia/Asphyxiation/Natural Causes | No |
| P1 | Sural nerve | M | 52 | Not Applicable | Type II Diabetes |
| P2 | Sural nerve | M | 60 | Not Applicable | No |

**Table S2:** RNAscope probes.

| Probe | Gene Name | ACD Probe Catalog Number |
|---|---|---|
| *NTRK1* | Neurotrophic Receptor Tyrosine Kinase 1 | 402631 |
| *PRPH* | Peripherin | 410231 |
| *SCN9A* | Sodium Voltage-Gated Channel Alpha Subunit 9; Nav1.7 | 562251 |
| *SCN10A* | Sodium Voltage-Gated Channel Alpha Subunit 10; Nav1.8 | 406291 |
| *SOX10* | SRY-Box Transcription Factor 10 | 484121 |
| *TRPV1* | Transient Receptor Potential Cation Channel Subfamily V Member 1 | 415381 |

**Supplementary files:**

**Suppl. File S1. (separate Excel file) Patient information**. Details on age, sex, diabetes, surgery procedure, related medical history, specimen collected, nerve morphology, sequencing and RNAscope.

**Suppl. File S2. (separate Excel file)** TPM values for tibial and sural sequencing data.

**Suppl. File S3. (separate Excel file)** Results of statistical analysis for sural versus tibial.

**Suppl. File S4. (separate Excel file)** Gene enrichment analysis for genes upregulated in sural nerves.

**Suppl. File S5. (separate Excel file)** Gene enrichment analysis for genes upregulated in tibial nerves.

**Suppl. File S6. (separate Excel file)** Results of statistical analysis for severe versus moderate axonal loss sural nerves.

**Suppl. File S7. (separate Excel file)** Gene enrichment analysis for genes differential expressed between severe versus moderate axonal loss sural nerves.

**Suppl. File S8. (separate Excel file)** Proteomics data. RNA-binding proteins (RBPs) detected in human DRG and sciatic nerves using Somascan assay.
